# Circulating DNA reveals nucleosome occupancy patterns that are associated with nucleosome-DNA affinity and are affected in cancer

**DOI:** 10.1101/2025.10.08.681110

**Authors:** Marianne Richaud, Ekaterina Pisareva, Paul Burgat, Alain R. Thierry, Jacques Colinge

**Author notes:** Correspondence should be addressed to Jacques Colinge or Alain Thierry.

## Abstract

The study of cell-free circulating DNA (cirDNA) fragments (fragmentomics) from liquid biopsies has received increasing attention. By constructing an atlas of these well-positioned nucleosomes, which we called WPNA, we found that their occupancy was associated with histone-DNA affinity, as evidenced by codon usage bias and differences in cirDNA fragment sizes. Moreover, WPNA nucleosome occupancy was different in healthy and cancer samples, thus allowing developing a high-performance machine learning approach for cancer detection (specificity and sensitivity >0.95 for seven cancer types). Cancer influenced WPNA nucleosome occupancy in a global manner, although distinct cancer types retained specific features. WPNA nucleosome occupancy at transcription factor binding sites revealed shared, pan-cancer regulation of transcriptional programs involved in hematopoietic cell differentiation and neutrophil biology, the main cirDNA sources. This work provides new fundamental insights into cirDNA and DNA sequence using cirDNA as a physical readout. It also bares translational significance by disclosing a new high-performance strategy for cancer detection from liquid biopsies.

## INTRODUCTION

Cell-free circulating DNA (cirDNA) can be passively (necrosis, apoptosis, phagocytosis) or actively released from a multitude of tissues and cell types, but mainly from hematopoietic cells (1–3). Therefore, it may convey information on the whole organism and constitutes a rich source of biomarkers. Landmark applications include non-invasive prenatal genetic tests, based on fetal cirDNA present in the mother’s blood (4), and cancer mutation detection in cirDNA released by tumors (5).

Upon release and in the blood circulation, cirDNA is actively degraded by different enzymes (6). The cirDNA fragments observed in blood are partially protected against these enzymes by DNA-binding proteins, such as histones or transcription factors (TFs) (7, 8). Sequencing of cirDNA fragments and alignment against the reference human genome determines a coverage at the base pair resolution and reveals stable nucleosome and TF occupancy *loci* (3, 9). Moreover, recent studies demonstrated that cirDNA methylation status brings additional information and allows deconvoluting the cells and tissues of origin (10, 11).

Fragmentomics is the analysis of cirDNA fragment characteristics, such as their size or nucleotide sequences at their extremities (end motifs). The cirDNA fragment size distribution is highly reproducible in healthy individuals, whereas it is altered in some diseases, thus providing a new biomarker type. For instance, in cancer, the size of the detected cirDNA fragments is typically shorter (7, 12), mostly due to the cancer-related enzymatic profile deregulation (13) and the overall increase in cirDNA release rather than due to direct cancer cell contributions (14). End motif frequencies also are altered in cancer and may provide additional biomarkers (13).

In this study, we bring new insights into the fundamental properties of cirDNA fragments. By developing a well-positioned nucleosome atlas (WPNA) that contains ∼5 million genomic positions. We found a relationship between genome coverage, fragment protection against enzyme degradation, and evolution of the genome sequence at these *loci*. Moreover, the genome coverage by cirDNA fragments at well-positioned nucleosomes was affected by cancer. This finding could be exploited to develop highly specific and sensitive cancer detection algorithms, including at early stages. Lastly, we investigated cirDNA genome coverage and its cancer-related changes globally and locally at transcription factor (TF) binding sites (TFBS).

## RESULTS

### Datasets and fundamental cirDNA fragment size patterns

The cirDNA fragment size distribution commonly observed in healthy individuals is described in Figure 1A. It displays a marked mode at 167-168 bp, which is the size of a nucleosome (145-147 bp) flanked by linker DNA protected by histone H1 (∼20 bp), *i*.*e*., a chromatosome. The distance between the small subpeaks, which are well visible on the left of the main peak, is typically 10.3 bp. This is the distance between two minor grooves along DNA wrapped around nucleosomes. This well-described phenomenon results from the fact that at these locations, the access of nucleases to nick DNA is easier.

**Figure 1.**
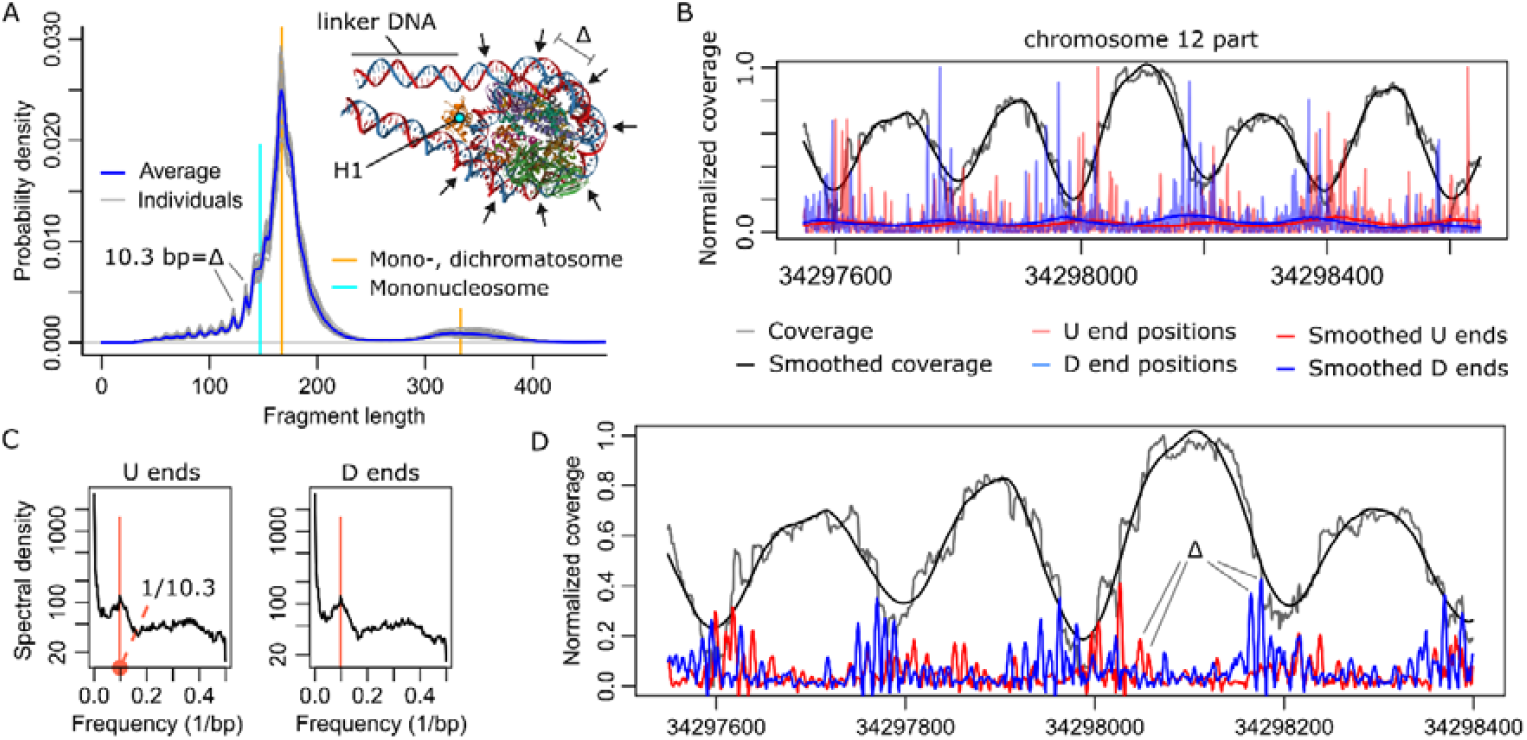
Features of cirDNA fragments. **(A)** Commonly observed cirDNA fragment size distribution in healthy individuals. The most represented size (mode) was at 167-168 bp, which is the size of a chromatosome (nucleosome with histone H1). The slight inflexion at 145-147 bp indicates a small population of nucleosomes without H1. Adjacent chromatosomes or other DNA-binding proteins protect larger DNA portions from enzymatic degradation and result in fragments larger 167 bp. Subpeaks are explained by facilitated access of enzymes at minor grooves of the nucleosome-wrapped DNA (small arrows, PDB structure 7K5X). The average distance Δ between successive minor grooves (10.3 bp) was the same as the distances between subpeaks. **(B)** Mapping of cirDNA fragments on the genome allowed computing cirDNA coverage. Representative example showing a portion of chr12. Fragments formed clear peaks at the positions of (stable) nucleosomes. Fragment ends were mapped more or less everywhere with a preference for linker DNA regions (smoothed red and blue curves, Savitzky-Golay filter). **(C)** The spectral density estimation (Welch method) of fragment end positions identified a clear peak at 1/10.3 = 1/Δ 1/bp. U ends = fragment ends at upstream genomic positions, D ends = downstream ends. **(D)** Reduced smoothing of fragment end positions (Savitzky-Golay filter) revealed a subtler structure with subpeaks every 10.3 bp in addition to the preference for linker DNA regions.

To identify novel properties of cirDNA fragments, we retrieved three cohorts of healthy individuals from FinaleDB (15) that were originally released by Cristiano, *et al*. (16), Jiang, *et al*. (17), and Sun, *et al*. (18). We named these cohorts after their first authors. They included 245, 32, and 13 individuals, respectively. As expected, the cirDNA fragment size distribution profiles in the three healthy cohorts were similar to what is described in Fig. 1A (Fig. S1). By modeling the overall shape of the cirDNA fragment size distribution profiles, we could subtract this shape to reveal the presence of subpeaks of increased fragmentation also after 168 bp (Fig. S2), indicative of continued DNA protection against nucleases by binding proteins. In the rest of the study, we used the largest dataset, *i*.*e*., Cristiano healthy cohort, for discovery and the other two datasets for occasional validations.

We then computed the genome coverage by the cirDNA fragments. Figure 1B shows a small part of chromosome 12 (chr12) where nucleosomes adopted stable positions that were revealed by well-separated peaks of coverage. As already reported (18), the upstream and downstream locations of cirDNA fragment ends (U ends and D ends, respectively) tended to occur preferentially at linker DNA positions, in agreement with the distribution mode at ∼167 bp (Fig. 1A). We performed spectral analysis of U and D end position occurrences in a large 65,636 bp window of chr12 that included the portion presented in Figure 1B. We found a peak frequency at 1/10.3 [1/bp] (Fig. 1C), suggesting the existence of a finer structure. Indeed, by selecting milder smoothing parameters, we revealed local, smaller peaks separated by 10.3 bp (Fig. 1D) that were reminiscent of the overall distribution structure.

### A notion of well-positioned nucleosome atlas and its occupancy

Different techniques have been proposed to putatively locate nucleosomes from cirDNA data. We based our approach on the window protection score (WPS) (3). For a given position *p*, the WPS is the ratio between the number of cirDNA fragments that cover an 80-bp window centered at *p and the num*ber of fragments that only intersect the window. As nucleosomes offer protection against nucleases, WPS peaks should reveal the position of nucleosomes. However, in genome regions where nucleosomes may adopt fuzzy positions, or where interindividual variability may occur, we found that WPS peaks alone might lead to ambiguous localization. Therefore, to locate well-positioned nucleosomes (19), we developed an algorithm in which WPS peaks above a minimum height were imposed additional conditions. We required sufficient separation between successive WPS peaks as well as the existence of local U and D end maxima up- and down-stream of a WPS peak (Figs. 2AB and Methods). In total, this algorithm identified 9,358,412 *bona fide* WPS peaks and estimated the corresponding cirDNA coverage peak diameters (or widths) (Fig. 2A). These diameters adopted a trimodal distribution (Fig. 2C). The main mode was close to 200 bp, which is a common nucleosome repeat length (NRL) in arrays of well-positioned nucleosomes (Fig. 1B). Larger diameters (>300 bp) were compatible with multi-nucleosomes, or overlapping fuzzy nucleosomes, while smaller structures (<147 bp) might be the product of TF-DNA complexes or other DNA-associated proteins. As our main objective was to describe well-positioned nucleosome locations, we only retained positions within the cirDNA coverage peak diameters of 147 and 300 bp. This constituted an *ad hoc atlas of 4*,971,768 well-positioned nucleosome positions (WPNA). We omitted chrY because we wanted sex-independent results. The number of nucleosome positions was proportional to the chromosome size (Fig. 2D).

**Figure 2.**
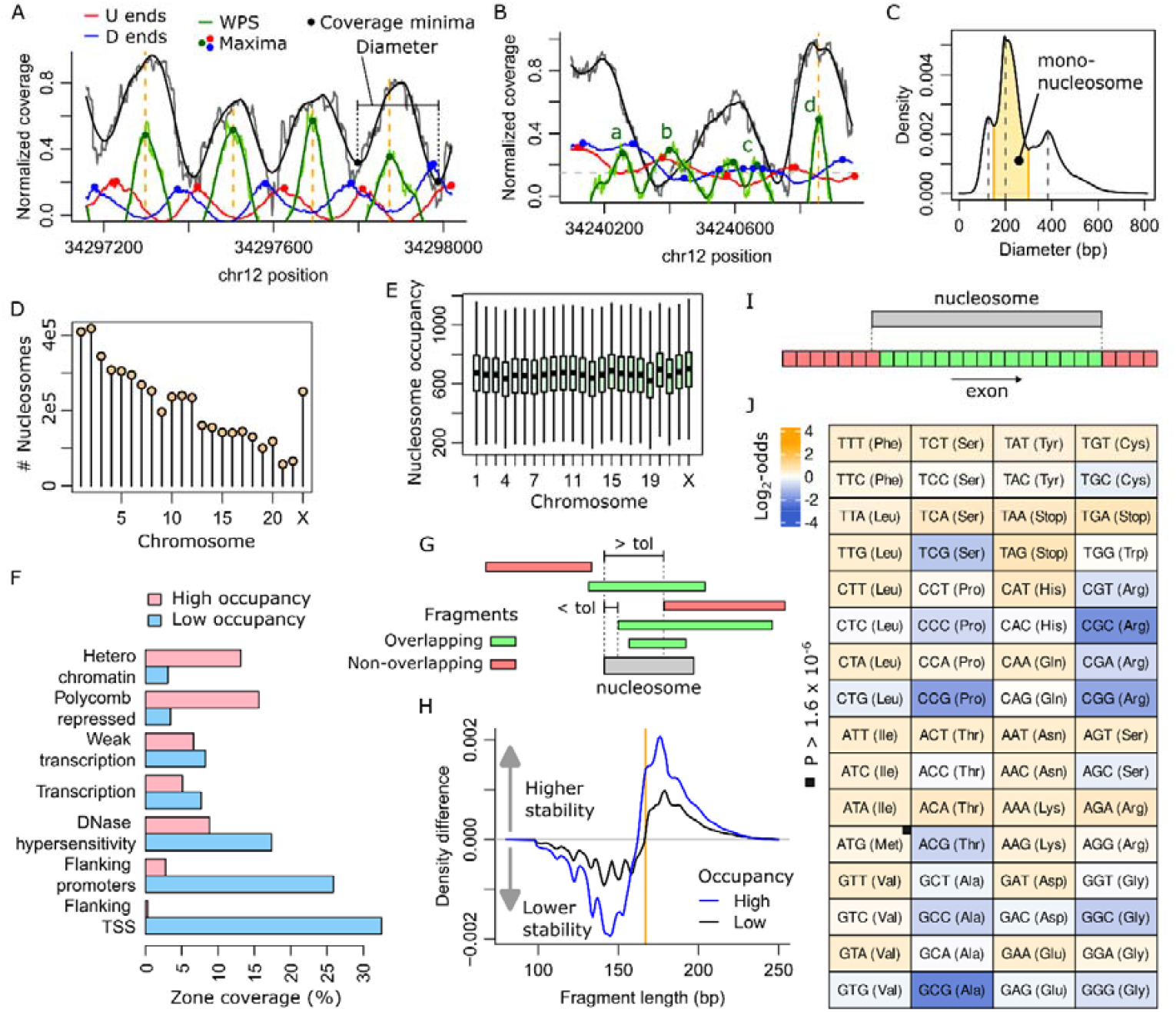
Nucleosome occupancy. **(A)** Illustration of the algorithm to define well-positioned nucleosome locations. WPS local maxima are candidate positions, provided they are well-separated from adjacent local maxima. Up- and down-stream local maxima in the smoothed density of U and D ends are also required. WPS, U and D ends are shown using an arbitrary scale to fit the normalized coverage scale. CirDNA coverage peak diameters were defined as the distance between the up- and down-stream coverage minima. **(B)** Example of rejected WPS maxima: (a) the D end maximum is too close; (b) the U end maximum is too close and the (smoothed) coverage too low (below the minimum dashed gray line); (c) the WPS peaks are too close to each other. Conversely, (d) fulfills our criteria and is accepted. **(C)** Diameters of the cirDNA coverage peaks. Diameters compatible with well-positioned nucleosomes, such as in (2A), are highlighted in yellow. The other WPS peaks were not included in the atlas. **(D)** Number of atlas nucleosomes *per* chromosome. **(E)** CirDNA coverage or occupancy at atlas positions. ChrX nucleosome occupancy was adjusted for sex. **(F)** Coverage (% of nucleotides) of each genome annotation by the top 20% (high) and bottom 20% (low) occupancy atlas nucleosomes. All differences were significant (, P < 10^-16^). **(G)** CirDNA fragments that overlap with a nucleosome completely or within a tolerance (tol) of 20 bp were considered as protected (green) by that nucleosome. **(H)** The sizes of cirDNA fragments protected by high-occupancy nucleosomes were compared with the sizes of fragments protected by the same number of randomly positioned nucleosomes. The relative size frequency differences (blue curve) showed higher prevalence of longer fragments. The same analysis for low-occupancy nucleosomes also showed an increase in size over random positions, but in much lesser proportion (black curve). **(I)** A nucleosome position compared with a protein coding gene exon. Only codons that were completely covered by the nucleosome were included in the statistics. **(J)** Log_2_-odds of the frequency of codon types that were covered by high-*versus* low-occupancy nucleosomes (*χ*^2^, P < 1.6 × 10^-6^, but for ATG where P = 0.075).

Although the cirDNA fragment genome coverage is commonly represented using normalized values between 0 and 1 (Figs. 1B and 2A), the actual coverage, *i*.*e*., *the numbe*r of cirDNA fragments at a given genomic position, can vary substantially. At WPNA nucleosome positions, this coverage defined nucleosome occupancy (20), which varied similarly in all chromosomes (Figs. 2E and S3). To assess the global coherence of the WPNA and WPNA nucleosome occupancy, we exploited generic annotations of genome features that were obtained by compiling epigenetic data for >100 cell types (21). We hypothesized that this aggregative approach was suitable for cirDNA fragments that originate from different cell types. We defined the top 20% and bottom 20% cirDNA-covered WPNA positions as high-occupancy and low-occupancy nucleosomes, respectively. We compared the proportions of their intersection with each zone of genome annotation (Figs. 2F and S4), which resulted in significant and expected biases that support the atlas relevance. For instance, high-occupancy nucleosomes were strongly enriched at heterochromatin, and mostly absent at flanking transcription starting sites (TSS) and promoters.

Next, we sought to relate nucleosome occupancy with cirDNA fragment properties. First, we defined a criterion to identify cirDNA fragments covered by a nucleosome, *i*.*e*., fragments that fully overlapped with a nucleosome or with a tolerance of 20 bp (Fig. 2G). By comparing the lengths of cfDNA fragments covered by high- and low-occupancy WPNA nucleosomes *versus* an identical number of random positions in the genome, we discovered that fragments protected by high-occupancy nucleosomes tended to be longer that those at low-occupancy nucleosomes (Fig. 2H). We confirmed this observation in the Sun and Jiang healthy cohorts using the WPNA generated using the Cristiano dataset (Fig. S5). As cirDNA fragments result from enzymatic attacks that are limited by the protection offered by histone binding, this finding suggested that high-occupancy WPNA nucleosomes correspond to DNA positions with increased affinity for histones or reduced enzymatic efficiency. These hypotheses imply local adaptation of the DNA sequence to adjust histone affinity or to change the frequency of motifs recognized by nucleases, and also to maintain the DNA ability to encode diverse features, such as genes. Therefore, we compared the frequencies of codon usage at high-*versus* low-occupancy WPNA nucleosomes that intersected their positions with known human gene exons (Fig. 2I). We found significant differences in the log-odds of codon frequencies at high-*versus* low-occupancy nucleosomes (Fig. 2J). All amino acids harbored both increased and decreased codon usage, which was compatible with the maintenance of the coding capacity. The only exception was alanine where all log-odds values were negative, although two were close to zero.

### Well-positioned nucleosome occupancy and cancer

It is well-known that cell epigenomes are substantially rearranged in cancer (24). Moreover, the global impact of cancer on the amount of cirDNA released in the circulation (14) could reflect cellular changes that might also modify the epigenomes of non-malignant cell populations. Therefore, we asked whether our WPNA could be used to probe nucleosome occupancy in individual patients. Each individual would then be associated with roughly 5 million quantitative features and machine learning (ML) approaches could be applied for cancer detection. First, we confirmed that although patchier, nucleosome occupancy could be extracted for individuals at the sequencing depth used by Cristiano *et al*. (Fig. 3A). Compared with the occupancy for the whole cohort (violet), occupancy for a given individual (black) varied for obvious biological and technical reasons (Fig. 3B). This variability needed to be taken into account by the ML procedure. As individual samples were not sequenced at constant depth and the cohort included women and men, we designed a normalization scheme before modeling WPNA nucleosome occupancy at the individual scale. We hypothesized that correct normalization combined with statistical testing at each WPNA position should lead to no significant differences. Specifically, to assess the sex bias, comparison of healthy women and healthy men should find no differences. By applying elementary normalization on the total occupancy over WPNA positions, we found many differential occupancies mostly on chrX, as expected (Fig. 3C). However, the PAR1 and PAR2 pseudo-autosomal regions yielded no difference, like in autosomes. Normalizing chrX separately corrected most of the problem, but introduced differences in PAR1 and PAR2. Finally, we opted for a normalization based on the total occupancy in autosomes, including the chrX PAR1/2 regions, and separately, chrX without PAR1/2. This led to acceptable false discovery rates (FDR) (Fig. 3C).

**Figure 3.**
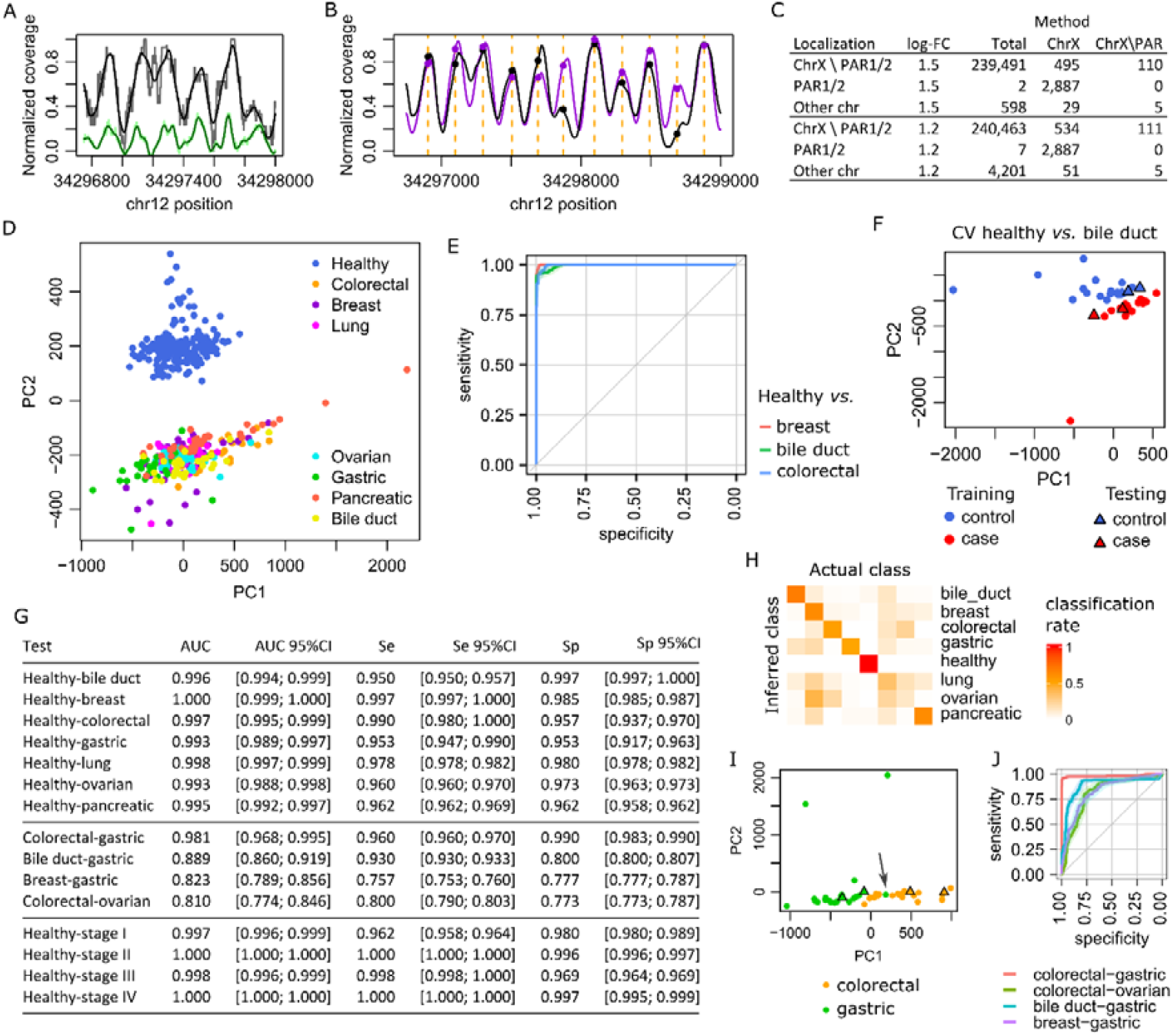
**(A)** Raw (gray) and smoothed (black) coverage of the healthy sample EE88155. The corresponding WPS curve (green) shows that cirDNA coverage peaks are clearly distinguishable. **(B)** Comparison of the smoothed individual coverage of sample EE88155 (black) and the whole Cristiano healthy cohort (violet). **(C)** Number of significantly different positional coverages between women and men (Wilcoxon, FDR < 0.01) at two minimum log_2_-fold change (FC) thresholds (in absolute value). For reference, the number of WPNA nucleosome positions on chrX was 480,452, including 8,387 in PAR1/2. **(D)** Two-dimensional PCA projection of healthy and cancer samples. **(E)** Representative receiver operating characteristics (ROC) curves for three Cristiano cancer type cohorts. **(F)** Illustration of our PCA-SVM classification algorithm and its cross-validation. **(G)** PCA-SVM classification performance (95%CI = 95% confidence interval). **(H)** Averaged confusion matrix of random forest classifiers illustrating how well distinct cancer types can be separated. Median overall accuracy = 0.77, 95%CI = [0.68; 0.82]. **(I)** Example of two cancer types that could be reasonably well separated. (J) ROC curves of easy and more difficult to separate cancer type pairs.

Then, we explored the nucleosome occupancy in cancer using the seven cancer cohorts (bile duct, breast, colorectal, gastric, lung, ovarian, pancreatic cancers) released by Cristiano *et al*. (16). Before comparing the Cristiano healthy and cancer cohorts, we used principal component analysis (PCA) to exclude potential strong outliers from each sample type. In each cohort, we found 1 or 2 outliers that were always the least cirDNA-covered samples (Fig. S6). The PCA projection of all the remaining samples resulted in very clear separation between healthy and cancer samples (Fig. 3D). Next, we designed an algorithm that combines PCA-based dimension reduction and support vector machines (SVM) for cancer detection using the cirDNA fragmentation data. This algorithm requires to embed the new samples to be tested with reference healthy and cancer samples in a common 2D projection. In this shared 2D space, the reference samples are used to build an SVM classifier, which is then applied to infer the status of the new samples. Due to the large difference in cohort sizes (244 healthy samples and 25 to 53 samples in function of the cancer type), and to avoid unbalanced learning, we repeated the 10-fold selection of random samples based on the smallest cohort (cancer) 10 times. Each time, a 2D PCA projection was computed and within this projection, 10-fold cross-validation (CV) was repeated 15 times to train/test the SVM models. We obtained highly sensitive and selective models (Fig. 3E). Figure 3F shows an example of PCA 2D projection with training and test data and Figure 3G summarizes the performance for cancer detection in individual samples for the seven cancer types. A simple linear model or linear discriminant analysis provided a similar performance, although slightly inferior (data not shown), indicating that SVM is just an option.

The PCA 2D projection (Fig. 3D) suggested that the separation of distinct tumor types was a more difficult ML task than cancer detection. Indeed, by keeping the first 20 dimensions in the PCA projection, we evaluated random forest (RF) models using 5-fold CV repeated 100 times. The average of the confusion matrices showed that some cancer types could be well separated (bile duct, breast, colorectal, gastric, pancreatic), but others were almost impossible to distinguish (ovarian, lung) (Fig. 3H and 3G). Overlap between training and test sets occurred even in the case of easily distinguishable tumor types, for instance colorectal and gastric cancer (Fig. 3I, and receiver operating characteristics (ROC) curves in Fig. 3J compared with Fig. 3E).

Lastly, to determine whether our approach could detect early stage tumors *versus* healthy controls, we pooled cancer samples (all seven cancer types) in function of their stage (I to IV). This task was again accomplished with very high sensitivity and selectivity by our algorithm (Fig. 3G). Unfortunately, our approach could not separate tumors in function of their stage (Fig. S7).

### Global properties of cirDNA genome coverage

As WPNA and the feature vectors associated with each individual were rather vast (4,971,768 measures of nucleosome occupancy), we asked whether we could select random nucleosomes from the WPNA to perform the classification with a much-reduced number of features. Namely, we randomly selected 20 sets of nucleosomes, and for each we performed the two embedded CV steps described above with 3 and 5 repetitions (instead of 10 and 15). In the case of the healthy *versus* pancreatic cancer samples, we used *N* = 10^5^,10^4^,10^3^, and we found that using only 10^5^ nucleosome positions instead of almost 5 million, gave the same performance (Fig. 4A). However, performance progressively decreased with *N* = 10^4^ and 10^3^. Breast cancer was easier to detect and we obtained a similar performance also with *N* = 10^4^ and 10^3^ (Fig. 4B). We obtained similar results for the other five cancer types, indicating that differences in nucleosome occupancy were rather global. Inspection of the cirDNA coverage for the healthy and cancer cohorts revealed many, rather small differences limited to a single or few consecutive nucleosome positions (Figs. 4C-D). Considering the WPNA-wide cirDNA coverage differences, we compared the distances in the dendrogram between the different cohorts after scaling the coverage over each chromosome in each cohort in the [0; 1] range (Fig. 4E). The colorectal and pancreatic as well as the breast and lung cancer samples constituted two rather close pairs. Ovarian cancer was closer to breast and lung cancer samples than the other cancer types, as previously shown by the confusion matrix in Figure 4h. Gastric cancer was the most different.

**Figure 4.**
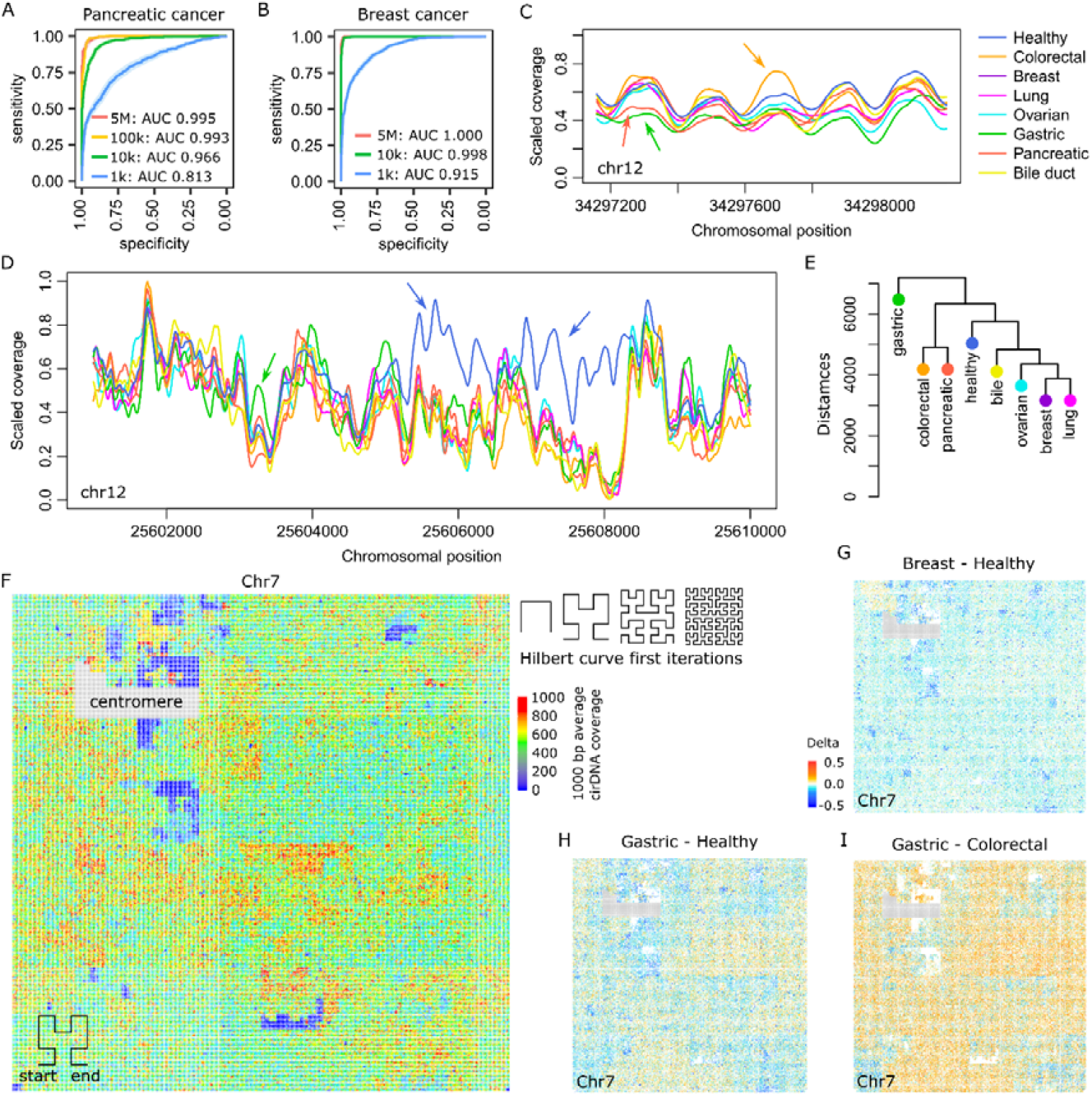
Global nucleosome occupancy. **(A)** ROC curves for the identification of pancreatic cancer versus healthy samples using the indicated number of nucleosomes randomly selected from the WPNA. (**B)** Same analysis for breast cancer. **(C)** Example of local variations in the different cohorts (arrows). Data were scaled in each cohort such that the maximum coverage per chromosome was 1. **(D)** Similar plot on a larger region to show one peak of higher coverage in gastric cancer (green arrow), and two sets of consecutive peaks of higher coverage in healthy samples (blue arrows). **(E)** Dendrogram showing the Euclidean distances of the indicated cohorts based on WPNA nucleosome occupancy differences. Occupancy values were scaled per chromosome (maximum = 1) in each cohort. **(F)** Two-dimensional mapping of chr7 cirDNA coverage (average every 1,000 bp). The principle of mapping linear information onto a unit square using iterations of the Hilbert curve is illustrated on the top right. Positions that were adjacent on the genome remained adjacent in the 2D image. Very low or absence of coverage occurred at locations away from the centromere. Similarly, high coverage was observed in rather specific areas. (G-I) 2D mapping of the WPNA nucleosome occupancy differences between healthy and breast cancer **(G)** and gastric cancer **(H)** samples and between gastric cancer and colorectal cancer samples **(I)**. To densify the difference plots and facilitate visualization, cirDNA peak diameter at each nucleosome position was multiplied by 4. Original plots in Figures S12-S14.

To represent the global organization of cirDNA coverage and WPNA nucleosome occupancy, we mapped such chromosomal linear quantities onto a unit square to facilitate their visualization. For this, we used a Hilbert space-filling curve (25), as exemplified for cirDNA coverage on chr7 in Figure 5F. We obtained similar qualitative results for the other chromosomes (Figs. S8 and S9 for chr12 and chrX, respectively): a locally patchy structure with pronounced global variations. Using the data of the Jiang and Sun healthy cohorts, we found that this structure was essentially preserved with some variations (Fig. S10). As expected from the correlation with genome annotations (Fig. 2F), mapping such annotations onto the unit square revealed positional patterns coherent with cirDNA coverage (Fig. S11). To complement the local analysis, we computed the differences between scaled nucleosome occupancy values. Illustrative examples in Figures 4G-H show more pronounced variations in gastric cancer than in breast cancer, as suggested by the dendrogram (Fig. 4E). Based on the latter dendrogram, we also computed differences between gastric and colorectal cancers and indeed observed larger values (Fig. 4I). Like for nucleosome occupancy only (Fig. 4F), differences were locally patchy, although they followed also global patterns that did not always match those of cirDNA coverage in the healthy cohort. This suggested the occurrence of changes that were not systematic. Lastly, it is important to note that nucleosome occupancy variations (Figs. 4G-I) remained modest in magnitude (see the color-coded scale), in agreement with the previous observation of small local variations. The patchy nature of differences at the global scale was also compatible with the finding that they were often limited to a single or few consecutive nucleosomes (Fig. 4C-D).

**Figure 5.**
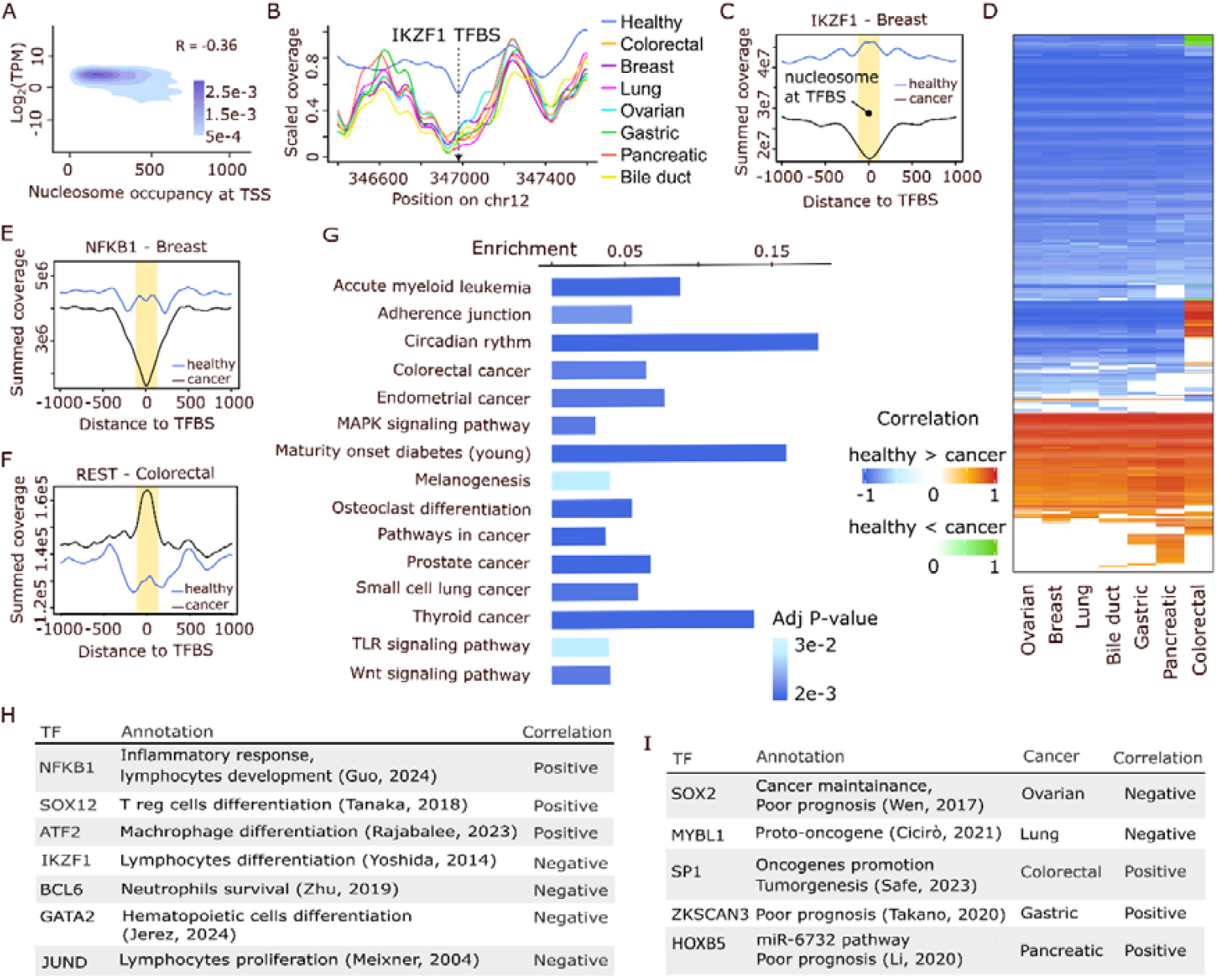
Changes at transcription factor (TF) binding sites. **(A)** Transcript expression levels in healthy hematopoietic cells in function of WPNA nucleosome occupancy at gene transcription start sites (TSS) in the Cristiano healthy cohort, and their Pearson correlation coefficient. **(B)** Total coverage in the heathy and cancer cohorts at a specific *locus* of chr12 that harbors the TF binding site (TFBS) of IKZF1. **(C)** Example of negative correlation between the healthy and breast cancer cohort total cirDNA coverage at the IKZF1 TFBS. Coverage is represented in a broad context of ± 1,000 bp, but the Pearson correlation coefficient was computed in the yellow window of ± 100 bp. **(D)** Heatmap showing the correlations of total cirDNA coverage in ± 100 bp windows harboring TFBSs in the seven cancer cohorts *versus* the healthy cohort. Two color scales were used to describe situations where coverage was higher and lower, respectively, in the healthy than cancer cohorts. **(E)** Same as in (6C) but for the NFKB1 TFBS that shows positive correlation between the healthy and breast cancer cohorts in the ± 100 bp window. **(F)** Same as in (6C), but for the REST TFBS that shows a positive correlation between the colorectal cancer and healthy cohorts, with higher coverage in the cancer than healthy samples. **(G)** KEGG pathways enrichment analysis of TFs for which nucleosome coverage at their TFBS was negatively correlated in all cancer cohorts *versus* healthy samples. **(H)** Examples of TFs with the same correlation patterns in all cancer types. **(I)** Examples of TFs displaying a cancer type-specific correlation pattern.

### Nucleosome occupancy and transcriptional regulation

Nucleosomes play an important role in transcription regulation. They can control DNA accessibility at specific positions, such as TSS and TFBS (9, 26). Therefore, nucleosome occupancy should reflect DNA accessibility at such loci in cirDNA-releasing cells. As previous studies reported that cirDNA mostly originates from hematopoietic cells, particularly leukocytes, in both healthy individuals and patients with cancer (11, 14). We used the Cristiano healthy cohort to correlate cirDNA coverage at the TSS of known genes with transcript abundance in hematopoietic cells (27). We found a negative correlation (Pearson correlation coefficient = −0.36) (Fig. 5A) in agreement with the fact that nucleosomes repress gene transcription.

The cirDNA fragment size range in the Cristiano healthy and cancer cohorts from FinaleDB (≥ 100 bp) was not adapted to find fragments protected by TFs, which are much shorter (9, 28). Therefore, by adapting published techniques (9, 29), we inferred DNA accessibility to TFs based on nucleosome occupancy at TFBSs. An example of TFBS with differential nucleosome occupancy is featured in Figure 5B where the total scaled occupancy was reduced in all seven cancer cohorts compared with the healthy cohort. To perform a systematic analysis, we downloaded the JASPAR database that provided the TFBSs for 644 human TFs (30) and selected only those that overlapped with WPNA nucleosome positions. Then, we compared (Wilcoxon test) the occupancy values in cancer (each type) and healthy samples at each TFBS-overlapping nucleosome to identify TFBSs at WPNA nucleosome positions with differential occupancy. We summed the cirDNA coverage at these TFBSs for each TF in healthy and cancer samples separately over ±100 bp windows (Fig. 5C). We imposed Pearson correlation coefficient values > 0.6 in absolute value (cirDNA coverage in healthy versus cancer samples) to avoid random signals (Fig. 5D). We found a limited number of prototypical patterns (TFs in Figures 5C and 5E-F). The two most recurrent configurations displayed higher cirDNA coverage in healthy than cancer samples and reduced nucleosome occupancy at the TFBSs. In the first case (Fig. 5C), the negative correlation indicated stronger transcriptional regulation in cancer samples. In the second case (Fig. 5E), the positive correlation indicated milder regulation in cancer samples that strengthened a reduction already present in healthy samples. The third configuration (Fig. 5F) was very rare and corresponded to increased TF binding repression in cancer samples (positive correlation).

Overall, we found 250 TFs with higher WPNA nucleosome occupancy in healthy samples and negative correlation in all cancer types and 97 TFs with higher WPNA nucleosome occupancy in healthy samples and positive correlation in all cancer types. The 250 negatively correlated, and thus more strongly regulated TFs were enriched in various KEGG (31) cancer pathways (Fig. 5G). The 97 positively correlated, and thus more mildly regulated TFs did not display any particular pathway enrichment. Inspection of the positively and negatively correlated TFs that were shared by all seven cancer types highlighted several TFs involved in hematopoietic cell differentiation: IKZF1 (32), NFKB1 (33), SOX12 (34), ATF2 (35), BCL6 (36), GATA2 (37), and JunD (38) (Fig. 5H).

Only few TFs adopted a cancer-specific nucleosome occupancy at their TFBS. Colorectal cancer had the largest number of specific TFs (n = 54), particularly many members of the SP-KLF family that have a role in this cancer regulation (39, 40). Other examples (Fig. 6I) were SOX2 (41) in ovarian cancer, SP1 (40) in colorectal cancer, MYBL1 (42) in lung cancer, HOXB5 (43) in pancreatic cancer, and ZKSCAN3 (44) in gastric cancer. Colorectal cancer was also the only cancer with TFBS positions more covered in cancer (14 TFs) than healthy samples. Some of these TFs had potential tumor suppressive functions, such as REST (Fig. 5F) (45) and EGR3 (46).

Neutrophil extracellular trap (NET) activation and release (NETosis) is an important source of cirDNA (47) and NF-kB was activated in neutrophils. IKZF1 is expressed in hematopoietic linages including neutrophils. GATA2 is required for survival and proliferation of multipotent progenitor cells that give rise to granulocytes, including neutrophils (patients with *GATA2* mutations often have present with neutropenia). SOX2 is a cytoplasmic DNA sensor in neutrophils. SP1 binds to GC-rich motifs in gene promoters, influencing the transcription of genes involved in neutrophil function. Namely, neutrophil elastase, an enzyme released by activated neutrophils, can stimulate the expression of genes, such as *MUC1*, by increasing SP1 binding to their promoters, indicating a role for SP1 in neutrophil-mediated gene activation. NF-κB signaling pathways cross talk with SP1 (48), and ZKSCAN3 negatively regulates these pathways in neutrophils. Besides these genes, we followed an unbiased approach to extract all genes defined in the KEGG NET formation pathway and then used the ChEA3 tool (49) to infer TFs that may regulate these genes. The intersection of the list provided by ChEA3 and our negatively correlated TFs was significant (P = 0.037, hypergeometric test).

## DISCUSSION

General properties of cirDNA fragments were searched in the large healthy cohort of 245 healthy individuals by Cristiano *et al*. (16). The cirDNA fragment size distribution in healthy individuals displayed the commonly observed pattern with a strong mode at 167-168 bp and subpeaks distanced by 10.3 bp on average. We found that these subpeaks also existed after the major mode at 167-168 bp. They are known to be caused by nuclease-facilitated access to protein-bound DNA at minor grooves (7, 8). In addition, the fragment U and D end distributions tended to concentrate in small intervals also separated by 10.3 bp on average. Moreover, the probabilities of observation at these small intervals augmented in the vicinity of nucleosome boundaries (18). Indeed, we observed both a local structure related to nucleosome positions and a microlocal structure related to nuclease activity and DNA nicks. As well-positioned nucleosome arrays typically adopt a ∼200-bp NLR (18), which is larger than a chromatosome (168 bp), we observed cirDNA fragment ends at positions that can be shifted by several multiples of 10.3 bp from the nucleosome boundaries. In a different context, relating DNA methylation and cirDNA fragmentation patterns, other authors observed periodic variations of fragment size probabilities relative to the nucleosome central position (50). This phenomenon is linked to the microlocal structure we evidenced in this work.

Nucleosome positioning is actively and accurately regulated in time and space through multiple mechanisms, such as DNA sequence, ATP-dependent nucleosome remodeling enzymes, histone and DNA modifications, poly(dA:dT) and poly(dG:dC) tracts, TFs, RNA polymerase II elongation and transcription, and non-histone DNA binding factors (19, 20). Moreover, nucleosomes assemble and disassemble constantly (22, 51). Therefore, we hypothesized that high-occupancy nucleosome positions would harbor distinct characteristics compared with low-occupancy positions. To address this question, we first designed a simple algorithm to assemble an *ad hoc* atlas of well-positioned nucleosomes that we called WPNA. We could assign a total of 4,971,768 nucleosomal locations to the WPNA built using the Cristiano healthy cohort. The number of WPNA nucleosomes *per* chromosome was proportional to the chromosome size, and occupancy varied identically in each chromosome in the [200; 1200] range (cohort total). High-occupancy WPNA nucleosomes were enriched in heterochromatin, and low-occupancy WPNA nucleosomes were enriched at gene-flanking promoters or TSS. These results confirmed the overall relevance of the WPNA as well as the existence of distinct properties in high- and low-occupancy WPNA nucleosomes. To identify physical differences, we exploited a naturally available readout: cirDNA fragment sizes. We discovered that fragments originating from high-occupancy WPNA nucleosomes were longer that those from low-occupancy WPNA nucleosomes. We could confirm this in two independent cohorts (17, 18). We hypothesized that such difference was related to variations in DNA-histone affinity. This implied that DNA locations harboring high-*versus* low-occupancy WPNA nucleosomes would have evolved slightly differently to adjust their affinity with histones and other proteins. However, at protein-coding genes, DNA is constrained to maintain its basic coding function. Therefore, we thought that evidence of DNA adaptation could be found in codon usage biases. Indeed, by computing the log-odds of codon usage in gene exons overlapping with high- and low-occupancy WPNA nucleosomes, we found clear differences.

Having demonstrated that WPNA nucleosomes associated with distinct DNA physical properties and genomic regions in function of their occupancy (high vs low), we asked whether occupancy was changed in cancer, although most cirDNA does not originate from tumor cells. Indeed, in cancer, malignant clone epigenetic profiles are profoundly affected (24) as well as cirDNA abundance (14), fragment size (7) and end motifs (13), to mention only the best described features. Cristiano *et al*. released seven cancer cohorts (bile duct, breast, colorectal, gastric, lung, ovarian, and pancreatic) in addition to the healthy individual cohort. From these cancer cohorts, we extracted the individual nucleosome occupancy levels for each patient at each WPNA position (4,971,768 measures *per* individual), and found that the PCA 2D-projection using these data could perfectly separate healthy individuals from patients with cancer. Therefore, we designed a new algorithm that combines PCA and ML and WPNA-related nucleosome occupancy data. This approach allowed discriminating between healthy controls and patients with cancer, including at early stages (AUC > 0.99, sensitivity > 0.95, specificity > 0.95 in all cases). The separation of distinct cancer types, a more difficult ML task, resulted in a heterogeneous performance. Patients with bile duct, breast, colorectal, gastric, and pancreatic cancer could be reasonably well separated from each other (median overall accuracy = 0.77, 95%CI = [0.68; 0.82]), but not patients with lung and ovarian cancer. In their study, Cristiano *et al*. achieved an overall accuracy of 0.6 for the same task (16). By exploiting other datasets and two to six different cancer types, other authors obtained overall accuracy values similar to ours (52–54). Our approach does not allow separating cancer stages within a dataset, but we are not aware of any method for doing this. For a different, although related purpose other researchers found a relation between cirDNA genome coverage and copy number variations (54).

As our PCA/ML algorithm could separate patients with cancer from healthy individuals very easily, cancer might have induced some global changes in nucleosome occupancy; however, very specific modifications might be limited in number and magnitude. This hypothesis was supported by the surprising discovery that randomly selecting only 10,000 to 100,000 nucleosomes from the almost 5 million contained in the WPNA was sufficient to obtain excellent cancer detection. To better understand this, we compared cirDNA genome coverage at the local and global scales. We observed rather modest local variations, limited to few consecutive nucleosomes. Conversely, global differences displayed a faint but clear structure. This global structure was cancer-specific and partially different from the basal cirDNA coverage, which was correlated with global genome properties such as closed/open chromatin and transcriptional activity.

Lastly, to complement this global analysis, we investigated the regulation of access to TFBS through WPNA nucleosome occupancy. We found that the TFBSs of 347 TFs were more available in all seven Cristiano cancer cohorts compared with the healthy cohort. Among other functions, these TFs are involved in cancer development, hematopoietic cell differentiation, and NETosis-related gene regulation. These functions reflect the main cirDNA sources (hematopoietic cells and NETs) as well as the contribution from cancer cells. A small number of TFs had access to their binding sites controlled by WPNA nucleosome occupancy in a cancer-specific manner. Colorectal cancer was the cancer type with the highest number of specifically regulated TFs (n = 54). The 14 TFBS with reduced accessibility in cancer were all detected in this cancer cohort and they included tumor suppressors. The rather homogeneous modulation of TFBS accessibility through well-positioned nucleosome occupancy agreed with the cirDNA global properties that varied little among cancer types. Therefore, we propose that cancer-specific nucleosome occupancy regulation does not occur predominantly at nucleosomes identified as well-positioned in healthy individuals. Their positioning might be deeply coded in our genome, such that even cancer has limited impact on them. Stronger changes might be observed at fuzzy nucleosomes, in cirDNA that directly binds to TFs and that was depleted in Cristiano’s cohorts (fragments were larger than 100 bp), and certainly at the more dynamic levels that are DNA and DNA-binding protein modifications. Furthermore, as we adopted a cohort view due to the limited individual sequencing depth, marked differences in patient subgroups might have been masked.

In conclusion, we identified new fragmentomic features related to well-positioned nucleosome occupancy that led to a highly sensitive and selective algorithm for cancer detection. We also exploited cirDNA as a physical readout to learn about the DNA sequence at well-positioned nucleosome *loci*.

## METHODS

### CirDNA sequence data

We downloaded cirDNA fragment sequence data from FinaleDB (15) as BED files representing the coordinates of cirDNA fragment alignment against the human genome assembly hg38. We used the data released by Cristiano *et al*. (16), Jiang *et al*. (17), and Sun *et al*. (18). We retained only fragments with alignment quality score > 30, and excluded non-chromosome genome assemblies (contigs).

### Software

For this study, a number of Python, R, and Perl scripts as well as C and C++ codes were written. These programs will be available at GitHub upon publication. Accordingly, here, we only describe the computation principles and not their implementation. Readers are invited to refer to software for such details. Note that the execution of some programs required significant computer memory and power and disk space. This cannot be done at the project-scale on a personal computer.

### Genome cirDNA coverage, fragment U/D ends

As previously described (3), we computed the number of cirDNA fragments that overlap with a given genome position by considering fragments between 120 and 180 bp in length. Such fragments represented the majority of fragments (Fig. 1a). We obtained the normalized coverage, as described in Figures 1B-D, by scaling the coverage between 0 and 1 in a 601 bp-wide sliding window. We obtained the smoothed coverage (actual or normalized values) by applying the Savitzky-Golay filter in 201 bp windows using 4^th^ degree polynomials.

We normalized cirDNA fragment U and D ends as described above and smoothed them by applying the Savitzky-Golay filter in 401 bp windows using 4^th^ degree polynomials (fig. 1b). For less smoothing (Fig. 1d), we applied the Savitzky-Golay filter in 13 bp windows using 4^th^ degree polynomials.

We estimated the spectral density of U/D end positions using the Welsh method (R library bspec).

### WPS computations and putative well-positioned nucleosome positions

We implemented *de novo* fast C++ code for WPS computation following the definition by Snyder *et al*. (27). We smoothed WPS values by applying the Savitzky-Golay filter in 101 bp windows using 4 ^th^ degree polynomials.

We identified putative well-positioned nucleosomes (before filtering based on the diameter) as local maxima of the smoothed WPS values > 0.15. We further imposed finding a local maximum of the smoothed fragment U end curve between 25 and 120 bp upstream. Similarly, we imposed a local maximum of the smoothed fragment D end curve between 25 and 120 bp downstream.

### Genome annotation intersection

We downloaded genomic annotations from Vu *et al*. (21). For the top and bottom 20% occupancy nucleosomes, using their respective diameters in the atlas, we computed the number of nucleotides that overlapped with each annotation zone. We divided this by the total length for each zone and expressed this ratio as a percentage.

### CirDNA fragment protection bias

Given a set of nucleosome positions, we first generated a matching set of randomized positions by adding to each original position a random number between −500,000 and 500,000 (uniform distribution), while avoiding to hit the centromere or to go beyond the chromosome ends. Then, we collected all cirDNA fragments in the chosen cohort, *e*.*g*., Cristiano healthy individuals, and identified those that overlapped with the nucleosome positions or their randomized counterparts. Each position (original or random) was associated with a window of ±100 bp, and we selected fragments completely included in a window (separately for original and random positions) as well as fragments that were not completely included, but with a non-included part ≤ 20 bp (tolerance).

To determine the bias of the original *versus* randomized selection, we computed the relative frequencies of each fragment length in the two selections separately, and then computed the difference between the original relative frequency and the randomized relative frequency.

### Codons usage bias

We downloaded the list of gene coding sequence positions in the human genome assembly hg38 from the Ensembl database (55) (Homo_sapiens.GRCh38.109.gtf). We filtered the coding sequence positions that intersected the top and bottom 20% occupancy nucleosomes. In both cases, we retrieved the nucleotide sequences, shifted the sequence starts to ensure that they started on an existing ORF, and split them into 3-bp triplets that corresponded to codons. We determined the relative frequencies of all codons for low- and high-occupancy nucleosome-covered gene sequence parts and this allowed us to compute log-odds (base 2 logarithm).

### Nucleosome DNA sequence signatures

We retrieved nucleotide sequences at *k*-stack positions for *k* = 0 (randomized 167-bp fragment positions), 1, 4 and 8. We also computed, as a preliminary step, 20-stacks to eliminate the fragments they contained as likely outsiders caused by repeat sequences. We counted nucleotide frequencies at each 167-bp fragment, from which we obtained PWMs. To compute the log-odds in the PWMs, we used 0.25 background probabilities for each of the four nucleotides (and base 2 logarithm).

We also computed the PWM for the dinucleotides AA/AT/TA/TT and CC/CG/GC/GG.

We carried out the spectral analysis with the Welch method as implemented in the R library bspec.

### Nucleosome occupancy and normalization in individual samples

We computed and smoothed the genome coverage by cirDNA fragments in individual samples as described above. We obtained the nucleosome occupancy at each WPNA nucleosome location by reading the smoothed coverage value of the individual sample.

We ignored chrY, and we normalized nucleosome occupancy at all the other chromosomes as follows. For each individual, we computed the total of the nucleosome occupancy values at all chromosomes, including the chrX PAR1 and PAR2 regions but not the non-pseudoautosomal part of chrX. Then, we divided the nucleosome occupancy values at these positions by the total and multiplied by the median of all total values. Likewise, we normalized chrX without the PAR1 and PAR2 regions by the individual corresponding total values of nucleosome occupancy and multiplied by the median of these totals.

### Principal component analyses

To efficiently compute PCAs of feature vectors with dimension close to 5 million required to use an iterative algorithm to avoid matrix products or inversions. We employed the optimized version of the Lanczos algorithm provided by the PRIMME library in R (56) to compute eigenvectors. With the *n* × k matrix *A* = (*a*_*i,j*_) containing normalized WPNA occupancy values (*n* values = #WPNA for *k* individuals or patients), we denote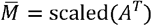, *i*.*e*., 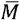 is equal to 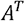 with each row scaled (subtract the row mean and divide by the row standard deviation). We used the R library matrixStats for faster mean and standard deviation computations. The correlation matrix is 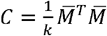 and the computation of principal components is equivalent to computing *C* eigenvectors. Indeed, given a vector *u* ∈ ℝ ^*n*^, we can write 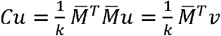, with 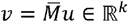. That is, we can chain two matrix-vector products and we do not even need to compute 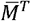 because 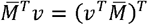. The function eigs_sym provided by the PRIMME library only requires to implement *Cu* to search for the *m* first eigenvectors. In practice, it happened that at a few WPNA positions, all patients had identical occupancy. We identified and discarded such positions beforehand to avoid dividing by a standard deviation = 0. We obtained the *m*-dimensional projections of the data in *A* by multiplying by the first *m* principal components (= eigenvectors here).

### Model construction, and AUC, sensitivity and specificity computations

We computed SVM models using the R library e1071. We computed the ROC curves and extracted the AUC values using the pROC package in R (57). To report balanced specificity (Sp) and sensitivity (Se) values, we determined for each ROC curve the point at the maximum distance from the diagonal. Elementary Euclidean distance computation shows that the distance between a ROC curve point (*x*_0_,*y*_0_) and the diagonal is given by 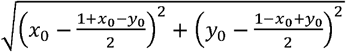. For a given CV, *i*.*e*., a ROC curve, we selected the farthest point that gave Se and Sp. To obtain 95% confidence intervals (95% CI), we applied a bootstrap (58) to the ROC curve points (R package boot, 100 re-samplings, balanced simulation, and CI from basic bootstrap). The pROC package computed directly the AUC 95% CI.

### Nucleosome occupancy at TSS

We downloaded the annotations of TSS positions from refTSS (59). We selected nucleosomes that intersected at least one TSS and determined their occupancy by summing over all Cristiano healthy samples at the center of each nucleosome. We obtained RNA-Seq data of healthy hematopoietic cells from the GEO dataset GSE74246. We converted counts to log_2_ TPM, and computed the mean nucleosome occupancy at all the TSS associated with each gene.

## ACKNOWLEDGMENTS

We thank Prof. Ya Ping Liu for developing FinaleDB and promptly answering our questions regarding data downloading and formats. His help was precious. MR was supported by a grant ANR RHU REVEAL to ART.

