## Supplemental information for "Circulating DNA reveals nucleosome occupancy patterns that are associated with nucleosome-DNA affinity and are affected in cancer"

### Supplementary Tables

**Table S1.** Definition of centromere boundaries used in all the computations.

| Chromosome | Centromere start (bp) | Centromere end (bp) |
|---|---|---|
| 1 | 125,100,000 | 143,200,000 |
| 2 | 93,900,000 | 96,000,000 |
| 3 | 90,900,000 | 94,000,000 |
| 4 | 48,200,000 | 51,800,000 |
| 5 | 46,100,000 | 51,400,000 |
| 6 | 58,400,000 | 60,300,000 |
| 7 | 58,100,000 | 61,000,000 |
| 8 | 43,900,000 | 46,100,000 |
| 9 | 38,500,000 | 69,000,000 |
| 10 | 38,500,000 | 42,200,000 |
| 11 | 50,700,000 | 54,700,000 |
| 12 | 34,600,000 | 37,400,000 |
| 13 | | 16,600,000 |
| 14 | | 20,200,000 |
| 15 | | 23,500,000 |
| 16 | 35,300,000 | 46,500,000 |
| 17 | 22,700,000 | 27,000,000 |
| 18 | 15,300,000 | 21,200,000 |
| 19 | 24,200,000 | 27,500,000 |
| 20 | 26,200,000 | 28,800,000 |
| 21 | | 13,100,000 |
| 22 | | 15,500,000 |
| X | 58,100,000 | 63,800,000 |

### Supplementary Figures

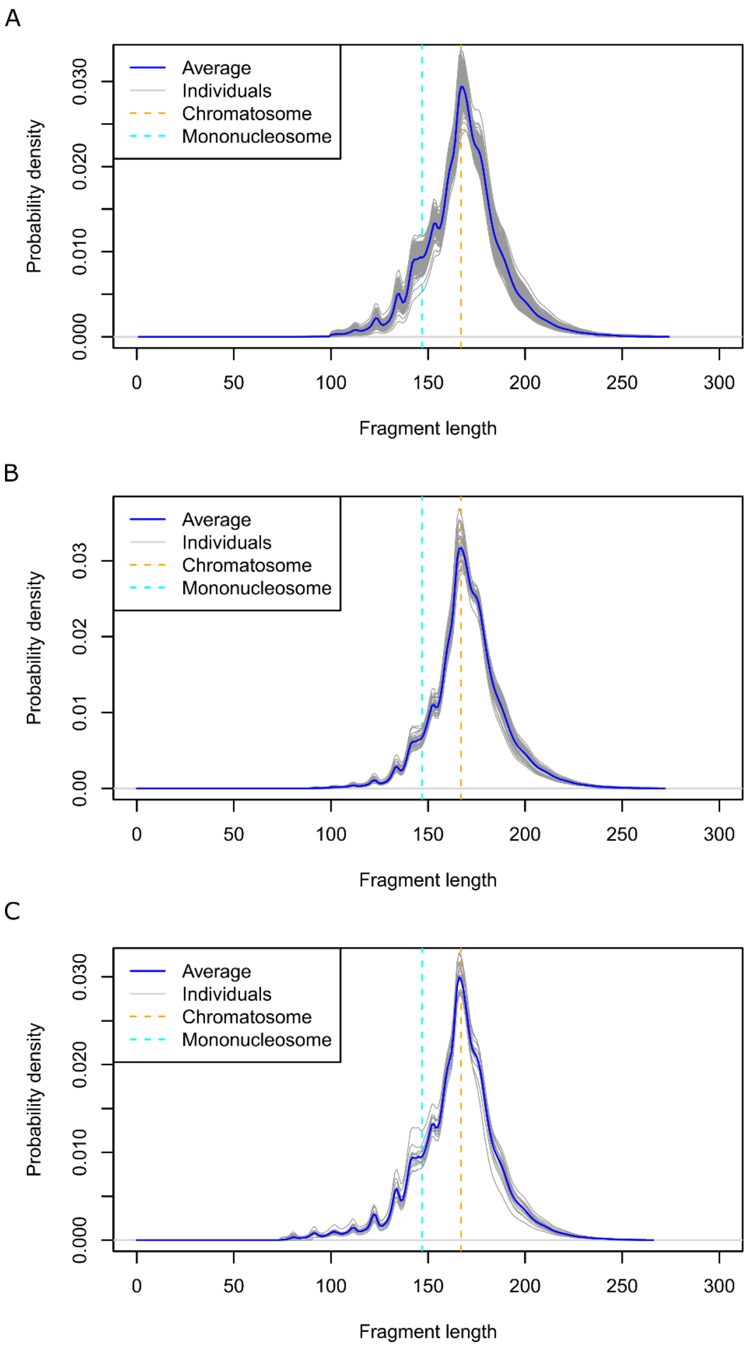


**Figure S1. CirDNA fragment size distribution.** (**A**) Cristiano, et al. data (2019) as retrieved from FinaleDB, $n$ = 245 healthy individuals. (**B**) Jiang, et al. data (2015) as retrieved from FinaleDB, $n$ = 32 healthy individuals. (**C**) Sun. et al. data (2019) as retrieved from FinaleDB, $n$ = 13 healthy individuals.

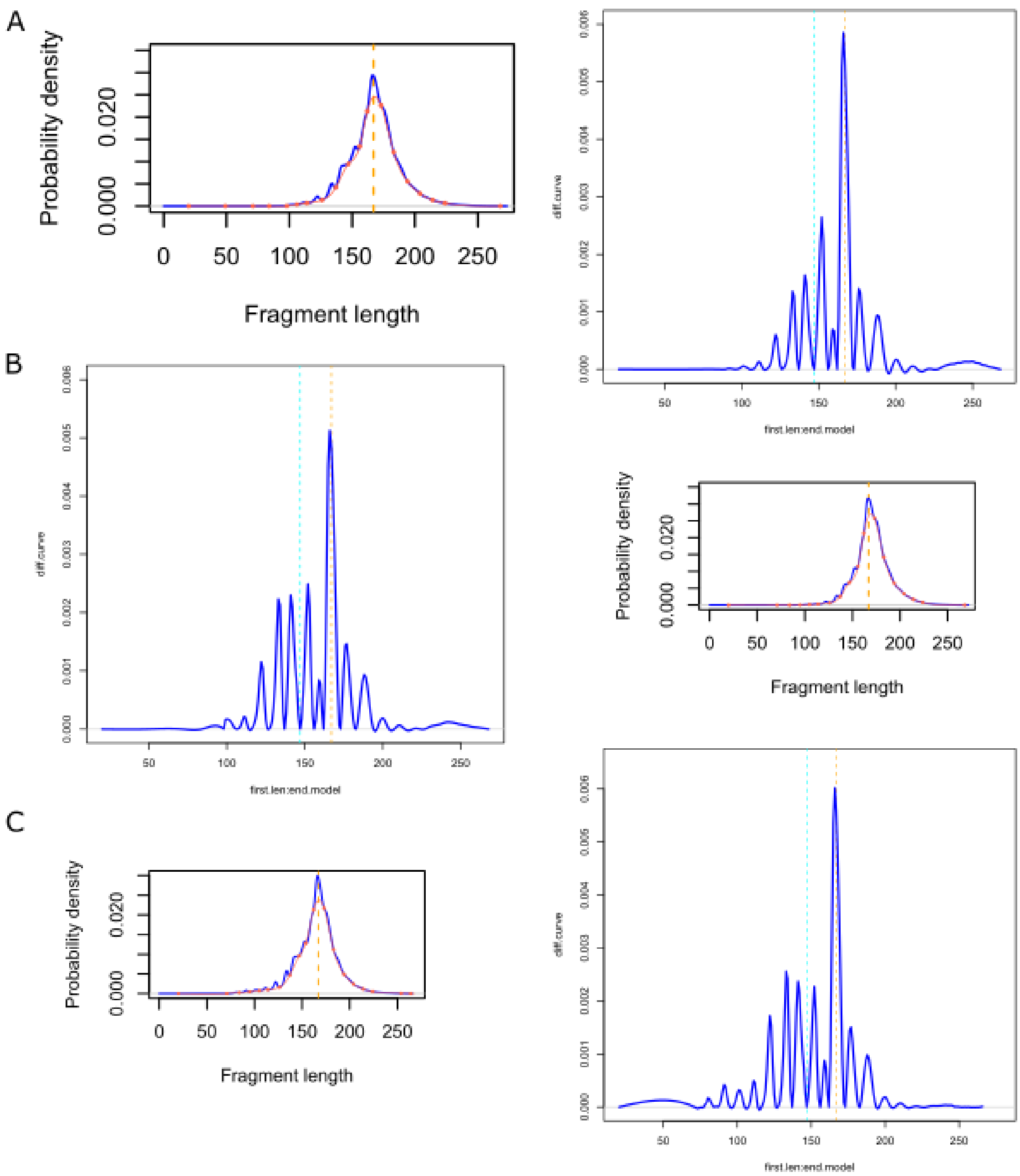


**Figure S2.** (**A**) Cristiano healthy cohort. Left, fragment distribution with a spline model to subtract the general shape of the distribution. Right, the subtraction reveals the small individual probability peaks, including on the right of the mode at 167-168 bp. Similar results were obtained for the (**B**) Jiang and (**C**) Sun healthy cohorts.

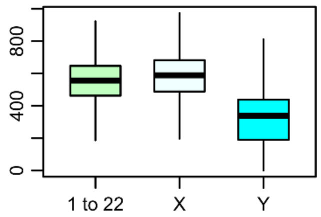


**Figure S3.** CirDNA fragment coverage of autosomes and gonosomes in Cristiano healthy cohort. Coverage is much lower for the chrY. Values were sex-corrected to make them comparable.

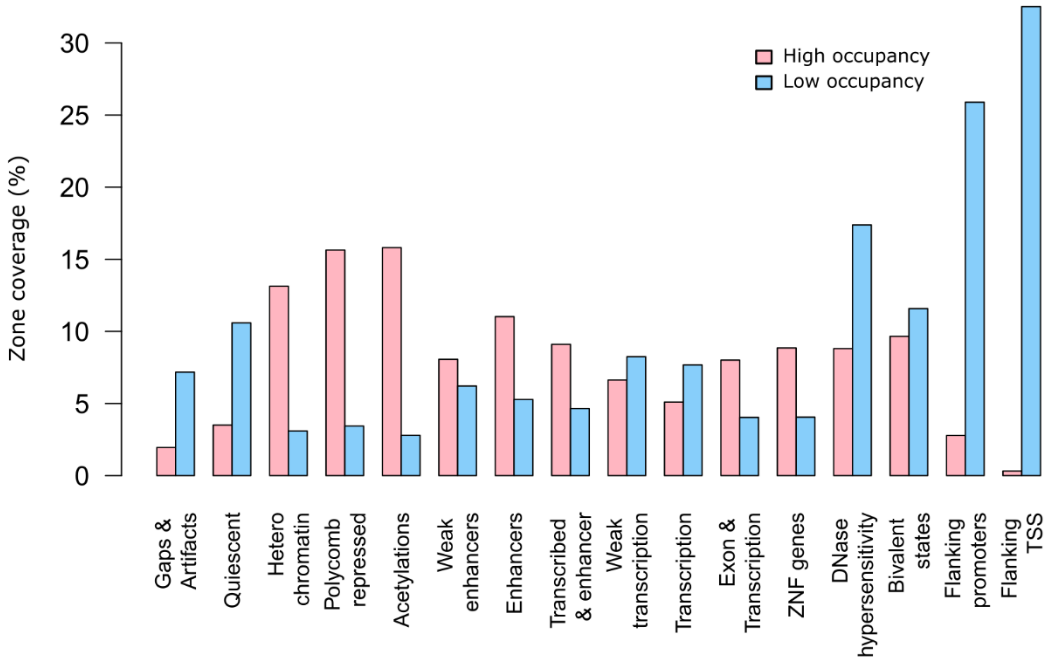


**Figure S4.** Full set of genome annotations.

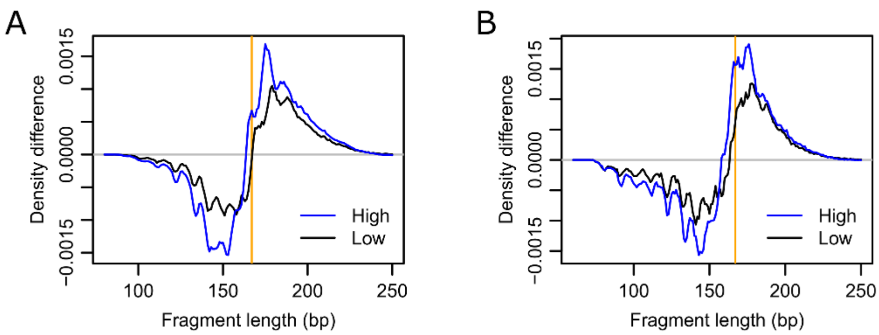


**Figure S5.** CirDNA protection by high- and low-occupancy WPNA nucleosomes *versus* random positions. (**A**) Jiang and (**B**) Sun healthy cohorts. The vertical orange line is at 167 bp. High/Low indicate top/bottom 20% cirDNA occupancy peaks.

A

| Type | Healthy | Colorectal | Pancreatic | Lung |
|---|---|---|---|---|
| Available | 244 | 25 | 34 | 38 |
| Outliers | 1 | 2 | 1 | 2 |
| Type | Breast | Ovarian | Bile duct | Gastric |
| Available | 53 | 27 | 25 | 25 |
| Outliers | 1 | 1 | 2 | 2 |

B

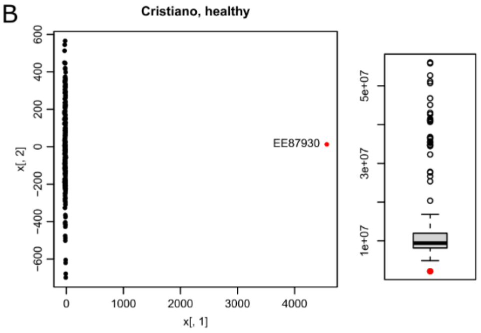


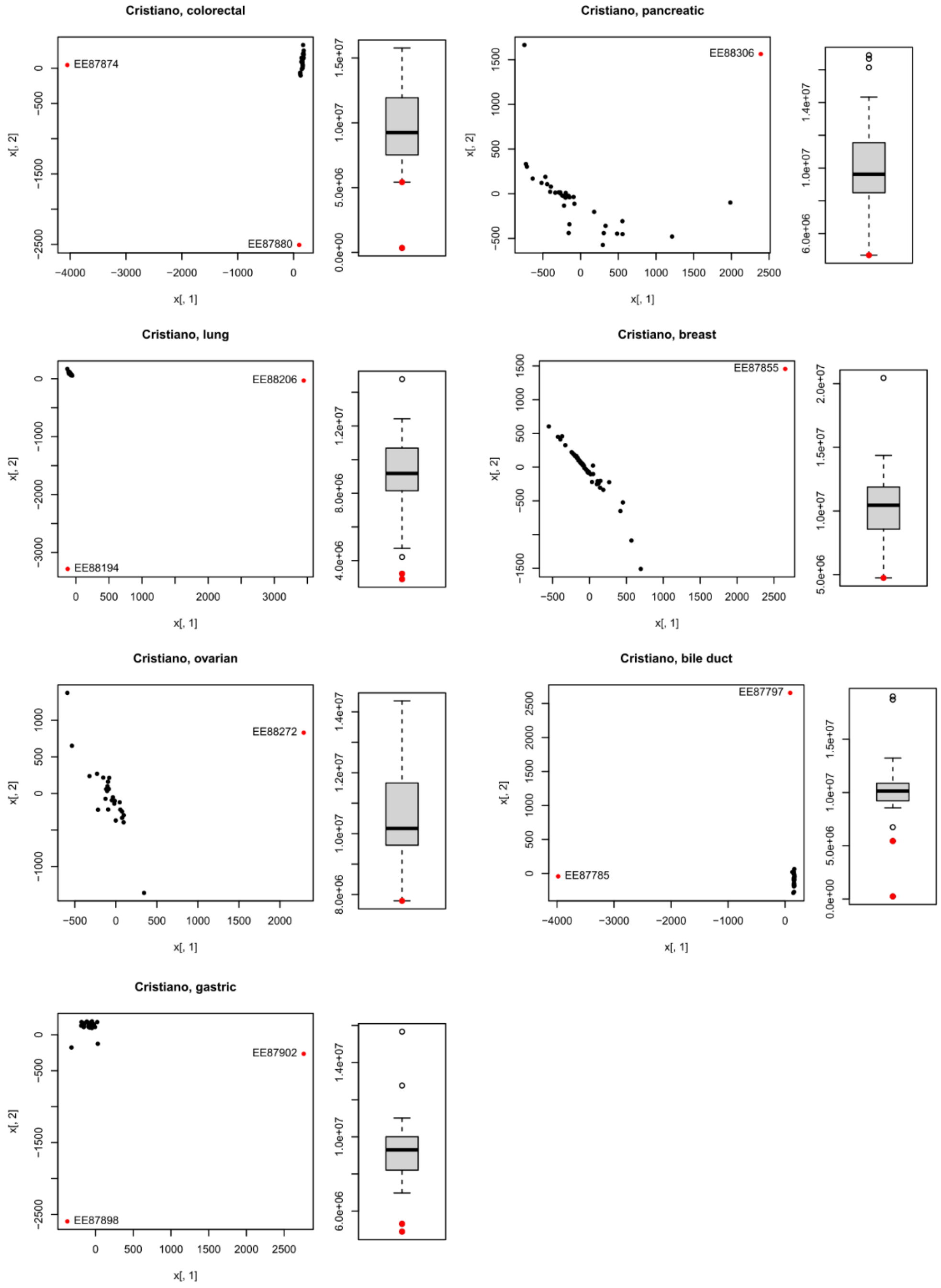


**Figure S6.** Outliers found in each Cristiano cohort. (**A**) Statistics. (**B**) PCA 2-dimensional projections followed by boxplots showing the total cirDNA coverage at WPNA positions in individual samples. Outliers are in red. Outliers were systematically the least cirDNA-covered samples.

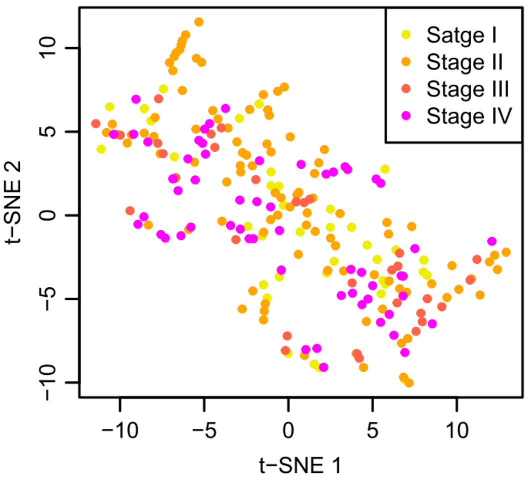


**Figure S7.** Reduction of the Cristiano cancer cohorts to 20 dimensions by PCA followed by t-SNE 2D projection. The four cancer stages could not be separated.

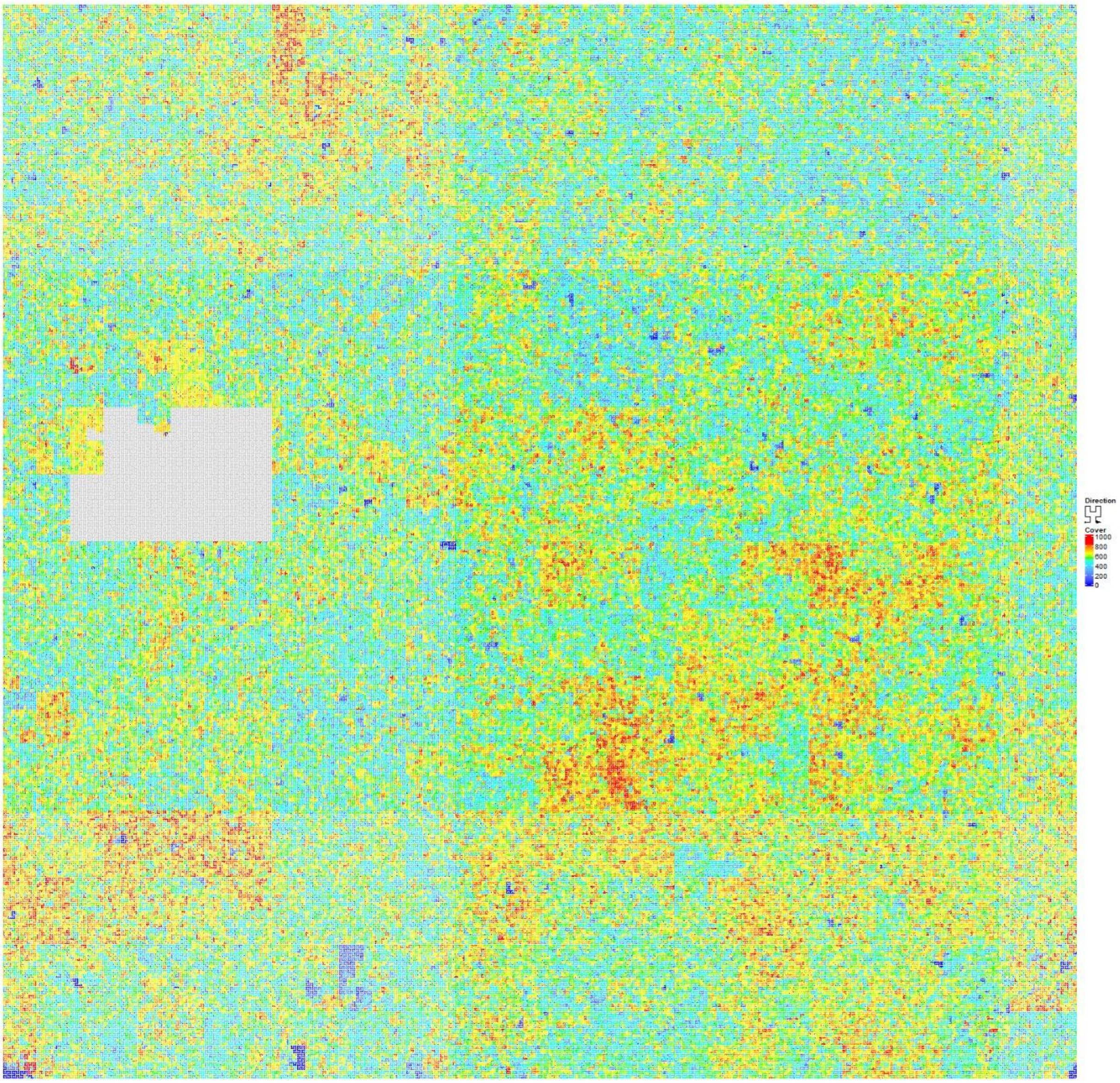


**Figure S8.** CirDNA coverage of chr12 (Cristiano).

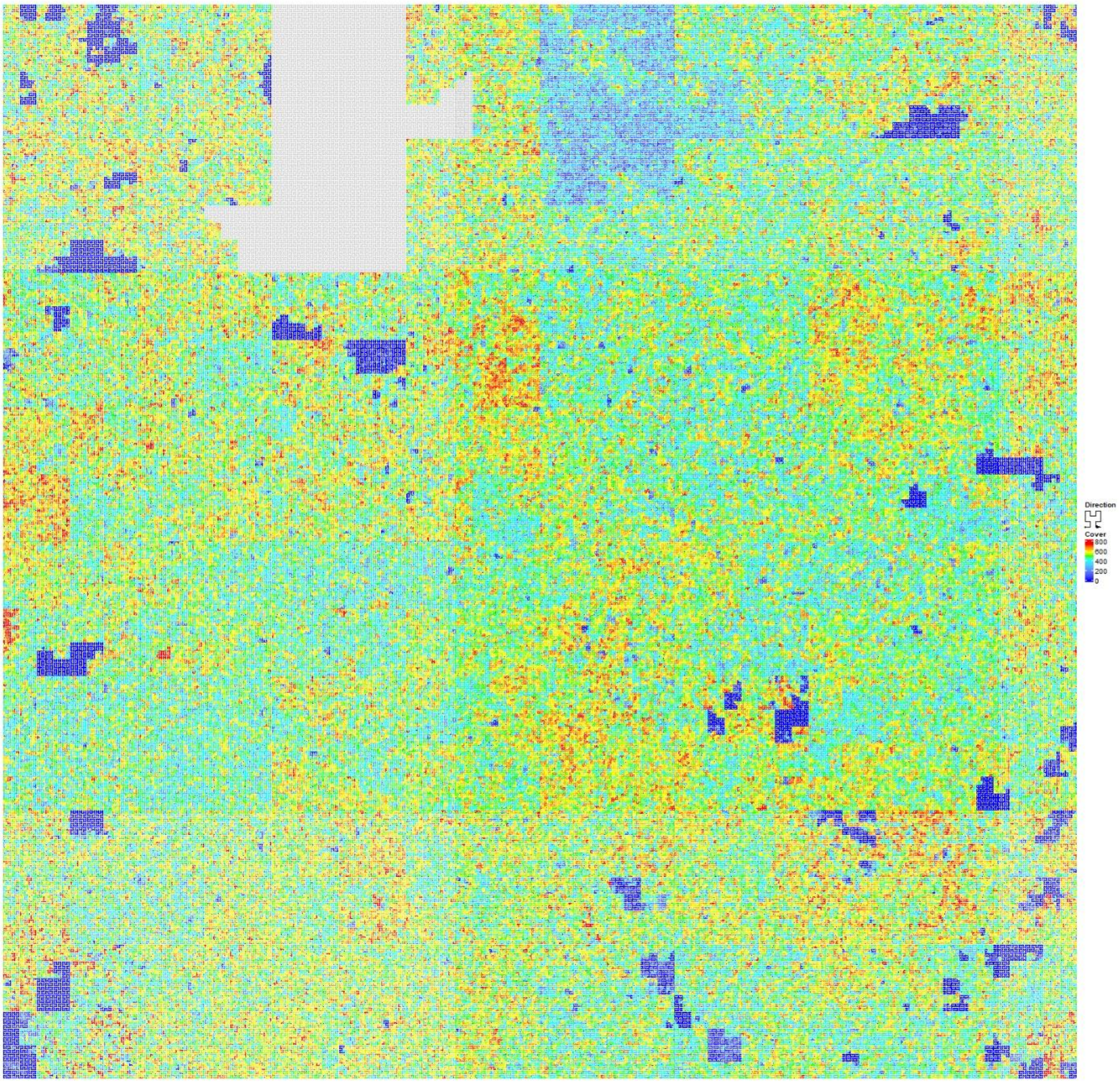


**Figure S9.** Cir DNA coverage of chrX (Cristiano).

A

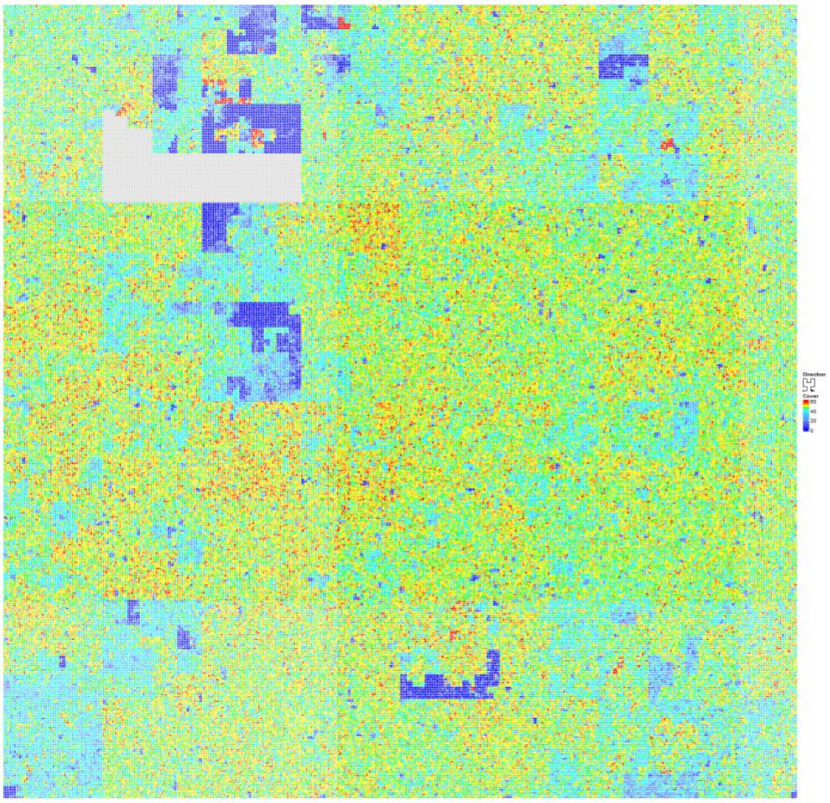


B

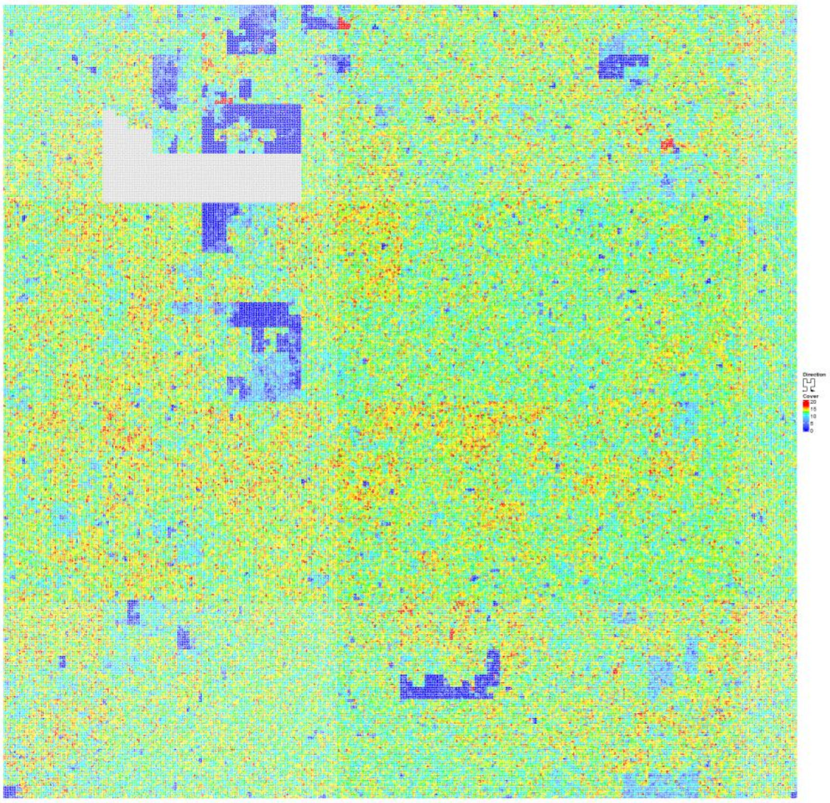


**Figure S10.** CirDNA coverage of chr7 in (**A**) Jiang data and (**B**) Sun data.

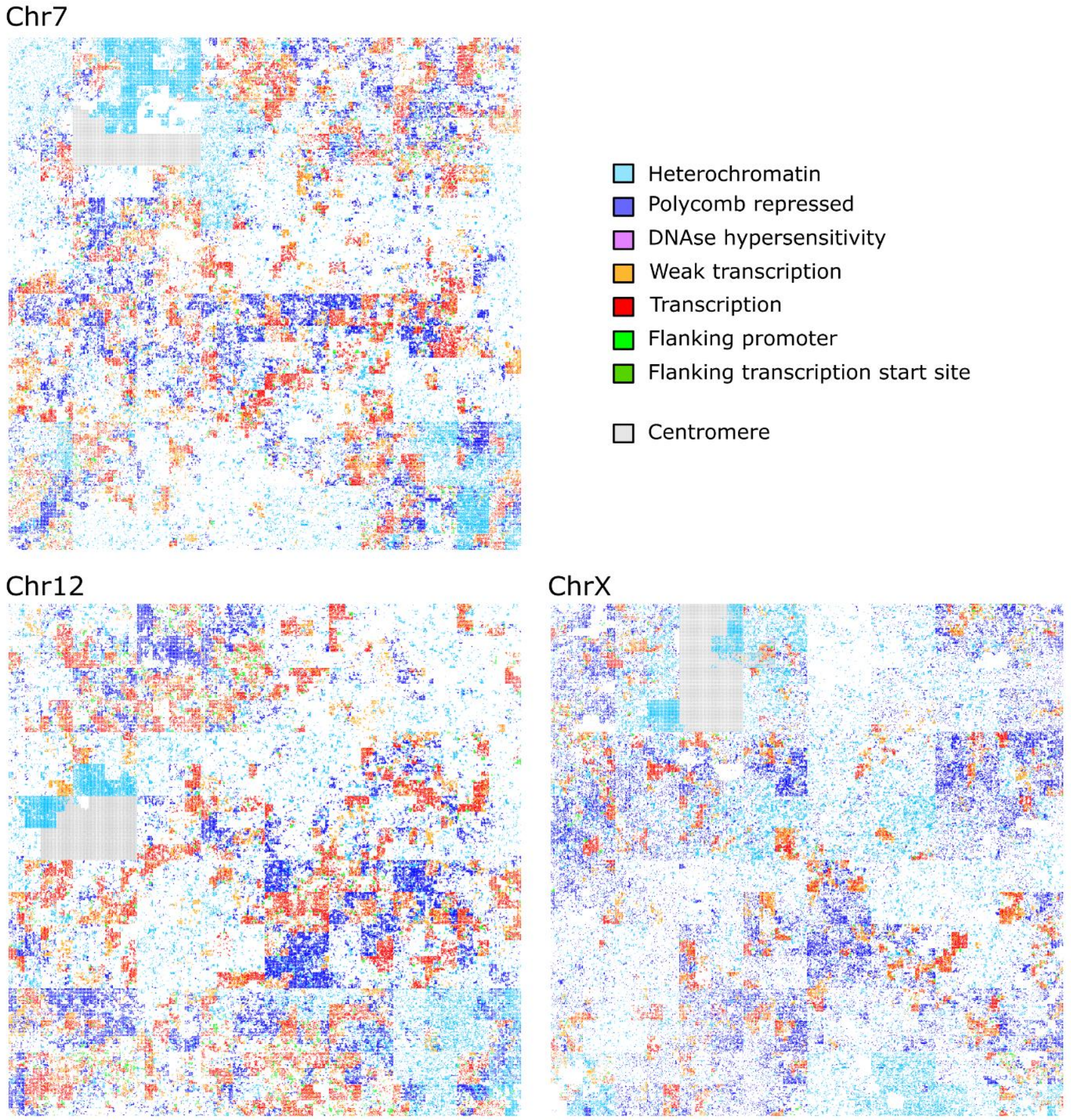


**Figure S11.** Mapping of chosen genome annotations. Annotations with blueish colors (heterochromatin, polycomb repressed) were associated with nucleosomes displaying higher occupancy, while the other annotations were associated with low-occupancy nucleosomes.

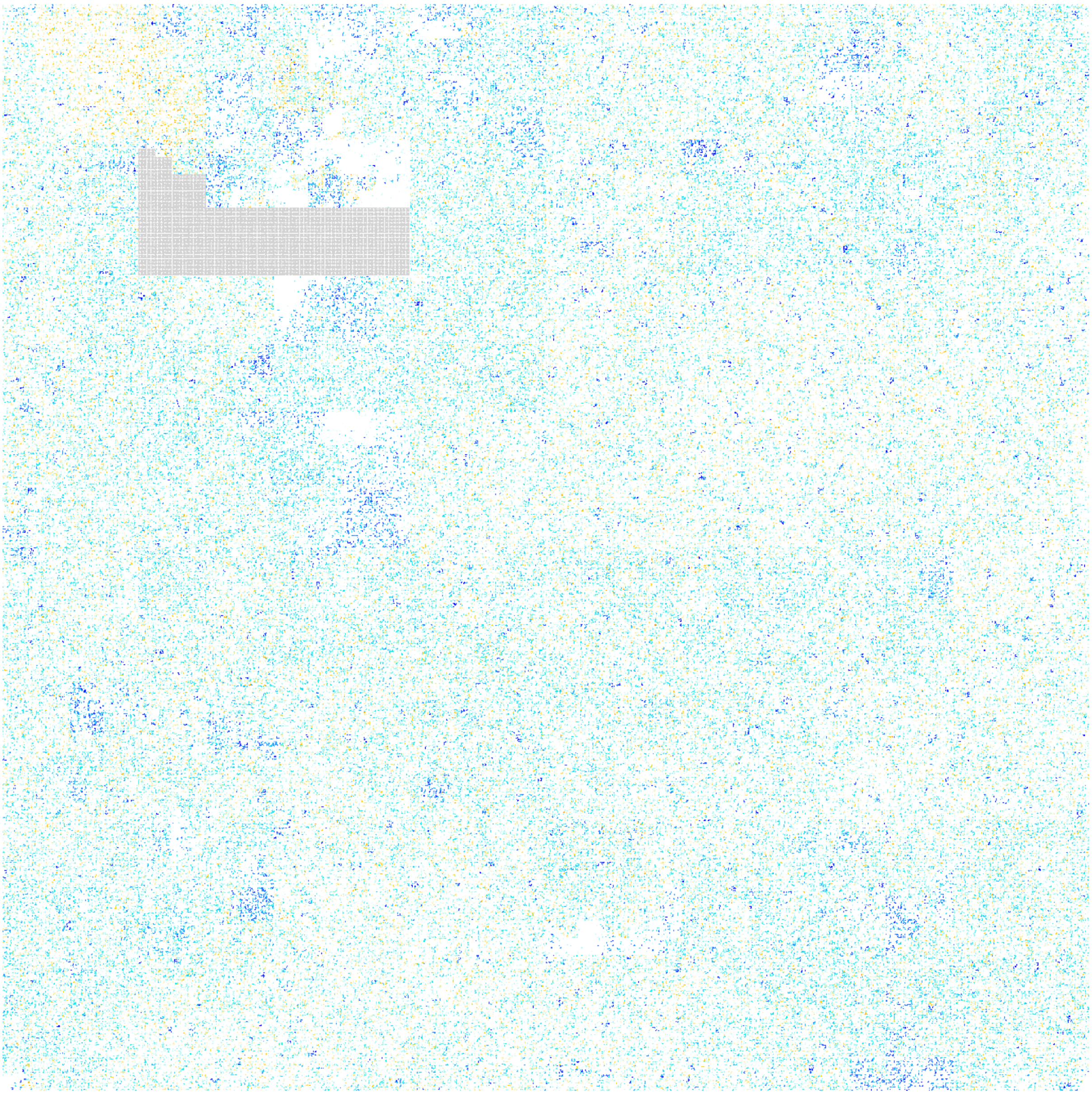

**Figure S12.** CirDNA coverage difference (after scaling in each cohort) between healthy and breast cancer cohorts.

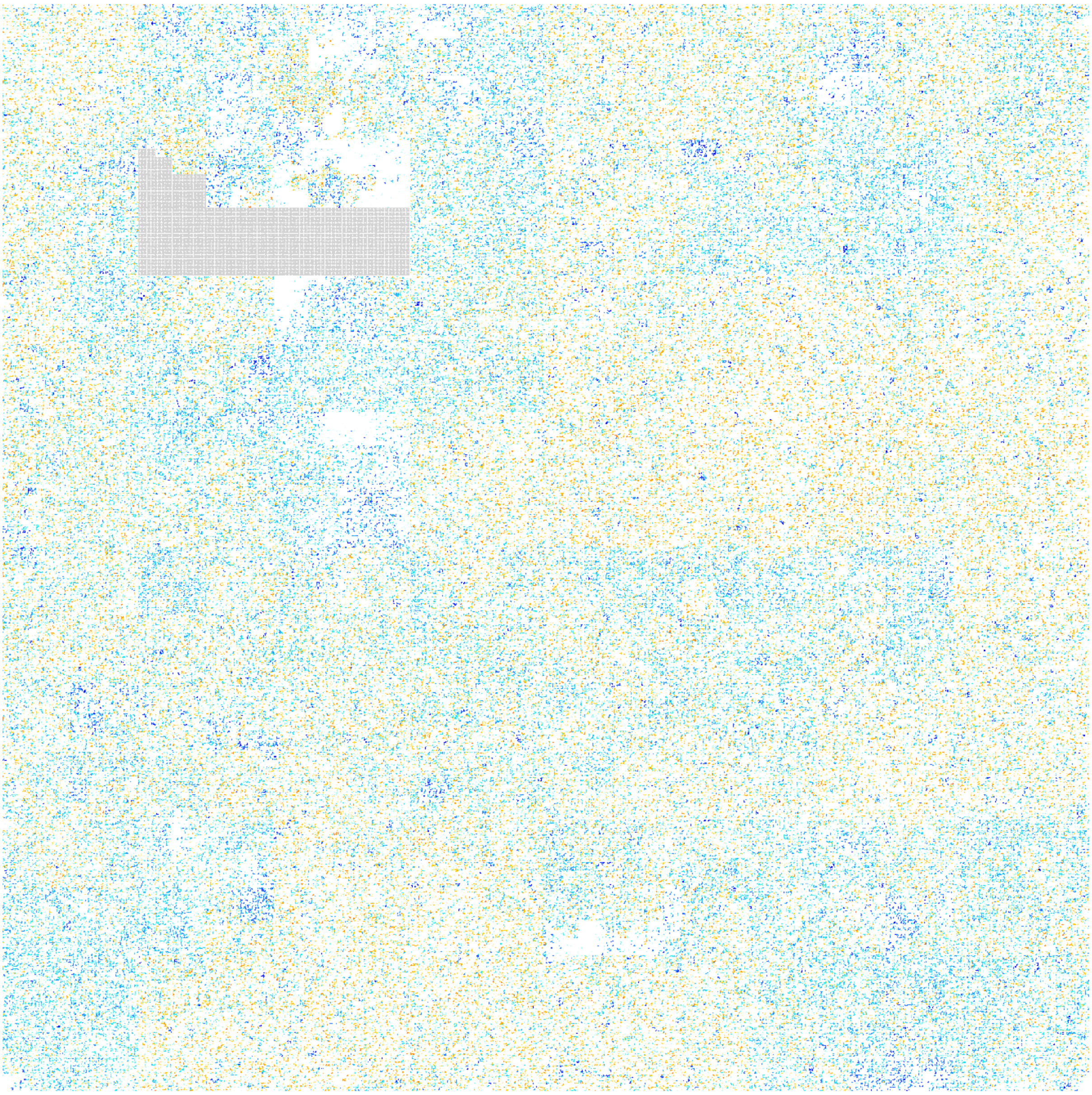

**Figure S13.** CirDNA coverage difference (after scaling in each cohort) between healthy and gastric cancer cohorts.

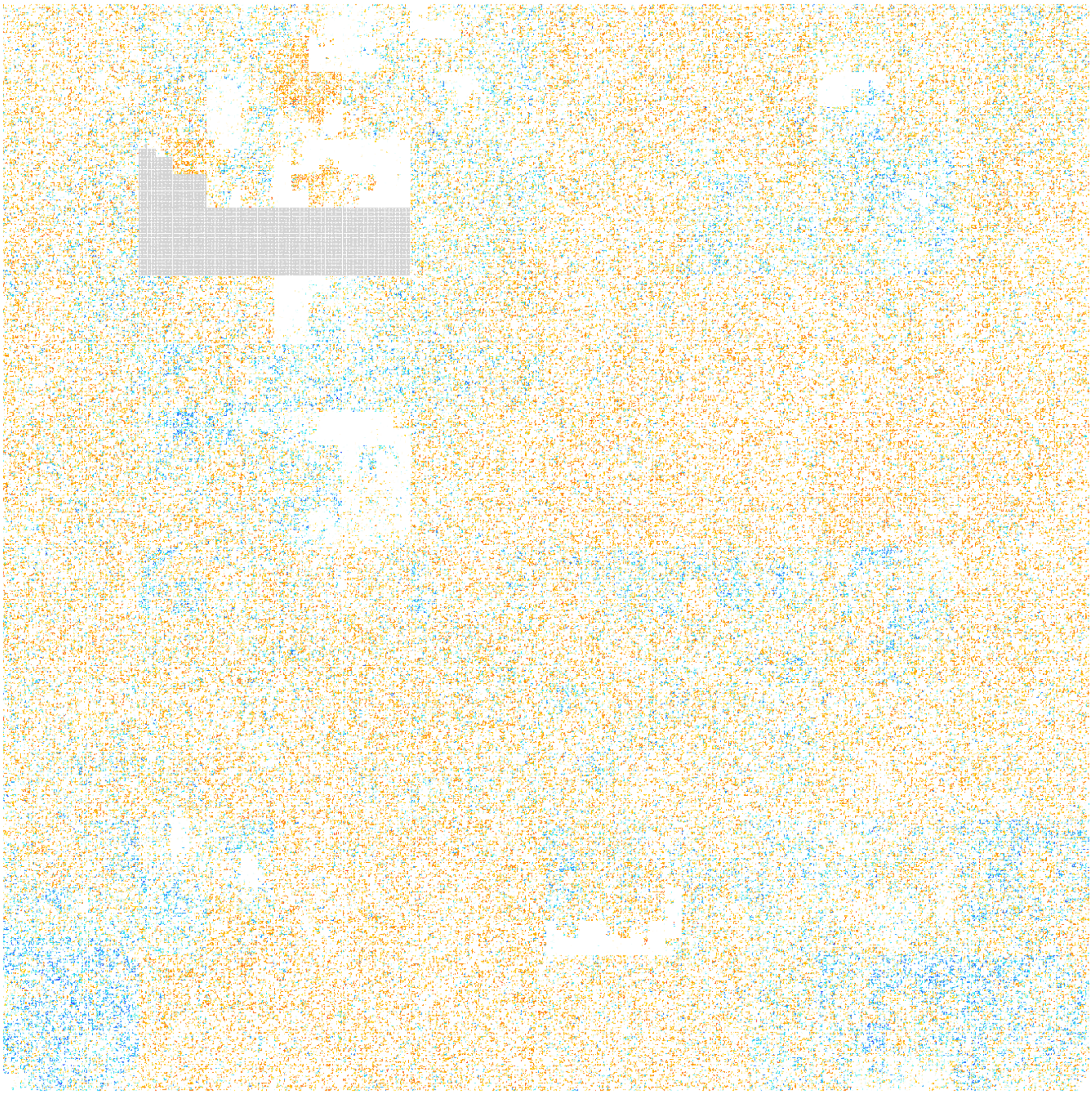

**Figure S14.** CirDNA coverage difference (after scaling in each cohort) between gastric and colorectal cancer cohorts.
